# The conserved centrosomal motif, γTuNA, forms a dimer that directly activates microtubule nucleation by the γ-tubulin ring complex (γTuRC)

**DOI:** 10.1101/2022.04.11.487887

**Authors:** Michael Rale, Brianna Romer, Brian P. Mahon, Sophie M. Travis, Sabine Petry

## Abstract

To establish the microtubule cytoskeleton, the cell must tightly regulate when and where microtubules are nucleated. This regulation involves controlling the initial nucleation template, the γ-tubulin ring complex (γTuRC). Although γTuRC is present throughout the cytoplasm, its activity is restricted to specific sites including the centrosome and Golgi. The well-conserved γ-tubulin nucleation activator (γTuNA) domain has been reported to increase the number of microtubules generated by γTuRCs. Here we utilize *Xenopus* egg extract and *in vitro* single molecule imaging assays to show that γTuNA activates microtubule nucleation in extract and directly activates γTuRC *in vitro*. Via mutation analysis, we find that γTuNA is an obligate dimer. Moreover, efficient dimerization as well as γTuNA’s L70, F75, and L77 residues are required for binding to and activation of γTuRC. Finally, we find that γTuNA’s activating effect opposes inhibitory regulation by stathmin. In sum, our study illuminates how γTuRC is controlled in space and time in order to build specific cytoskeletal structures.

## 3. Introduction

Microtubule (MT) assembly is a critical cellular process tightly regulated in both space and time. Spatiotemporal control of MT nucleation allows cells to use the same pool of soluble tubulin to generate different intracellular structures, from the interphase cytoskeletal transport network to the complex mitotic spindle. Yet, while the core MT nucleation machinery has been well characterized, how MT nucleation is regulated remains poorly understood.

The key MT nucleator is the γ-tubulin ring complex (γTuRC). γTuRC is a large, 2.2 MDa complex that forms an asymmetric ring of γ-tubulin subunits (Zheng et al., 1995; Moritz et al., 1998). This ring is thought to act as an initial template for the MT (Moritz et al., 2000). As α/β-tubulin subunits bind to the ring of γ-tubulin, they form the nucleus of a new MT, rapidly transitioning from nucleation toward the more favorable regime of MT polymerization (Jackson and Berkowitz, 1980; Mitchison and Krischner, 1984). *In vitro* studies with purified human and *Xenopus* γTuRCs have shown that these can indeed catalyze the nucleation of new MTs (Choi et al., 2010; Thawani et al., 2020; Liu et al., 2020). Recent studies have also shown that γTuRC acts synergistically with the MT polymerase, XMAP215/ch-TOG, to nucleate MTs (Thawani et al., 2018; Flor-Parra et al., 2018; Gunzelmann et al., 2018; B. King et al., 2020).

The centrosomal scaffold protein Cdk5rap2, which recruits γTuRC to the centrosome and Golgi (Anderson et al., 2003; Bond et al., 2005; Fong et al., 2008; Choi et al., 2010; Mennella et al., 2012; Lawo et al., 2012), has also been shown to increase γTuRC’s nucleation activity (Fong et al., 2008; Choi et al., 2010; Roubin et al., 2013). Previous domain-mapping studies found that the γ-tubulin nucleation activator (or γTuNA) sequence in Cdk5rap2’s N-terminus is critical to bind and activate γTuRC (Fig. 1A; Fong et al., 2008; Choi et al., 2010). Direct binding of γTuNA has been proposed to activate γTuRC, as addition of γTuNA increases γTuRC activity both *in vitro* and in cells (Choi et al., 2010). Prior work has also identified a key hydrophobic residue in γTuNA, F75, that is critical for binding and activation of γTuRC (Fong et al., 2008; Choi et al., 2010). The γTuNA domain is well-conserved across yeast, nematodes, flies, frogs, and humans (Samejima et al., 2010; Conduit et al., 2014; Feng et al., 2017; Fong et al., 2008; Choi et al., 2010), and γTuNA domains have been identified in related centrosomal and Golgi proteins such as myomegalin (Roubin et al., 2013). A bipartite version of γTuNA is also present in the microtubule branching factor TPX2 (Alfaro-Aco et al., 2017). Thus, understanding how the γTuNA domain interacts with γTuRC might bring insights into the regulation of MT assembly in a wide variety of organisms and contexts.

**Figure 1.**
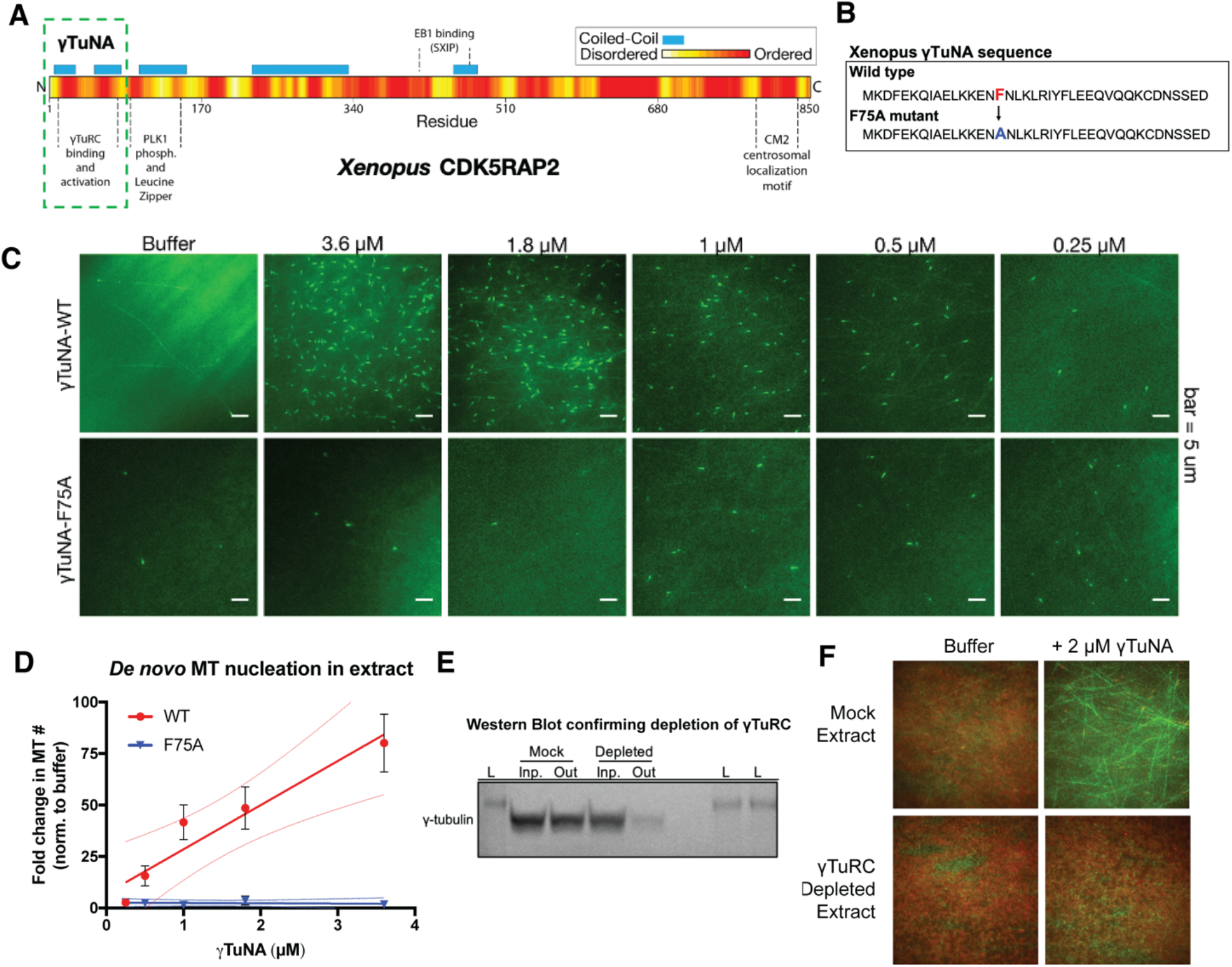
Cdk5rap2’s γTuNA domain increases MT nucleation in Xenopus egg extract and requires the universal MT template, the γ-tubulin ring complex (γTuRC). **A**) Schematic of *Xenopus* Cdk5rap2’s domains. The N-terminal γTuRC binding and activation domain, γTuNA, is highlighted in the green, dashed box. Predictions of disorder (PONDR-FIT) and coiled-coil regions (COILS) are shown in red and blue, respectively. **B**) Sequence of residues 60-98 (based on human numbering) for the wildtype and F75A mutant versions of *Xenopus* γTuNA. **C**) TIRF assay of MT nucleation in *Xenopus* egg extract. A titration series of wildtype or F75A versions of *Xenopus* γTuNA (Strep-His-γTuNA) were added to extract as shown. EB1-mCherry was used to mark growing MT plus-ends (pseudo-colored green in images). Bar = 5 μm. **D**) Quantification of the number of EB1 spots in C. The data were normalized by the buffer controls, and are shown as fold-changes. Black error bars are the standard deviation for four independent experiments. Thin colored lines on either side of the central trendline represent 95% confidence intervals. **E**) Western blot of γ-tubulin levels before and after mock-treatment or incubation with Strep-His-Halo-*Xenopus* γTuNA-coupled beads. After a single pulldown, the majority of γ-tubulin signal is lost. **F**) TIRF assay of mock- and γTuRC-depleted extract. Alexa-488 labeled tubulin (green) and EB1-mCherry (red) were used to visualize microtubules in extract with or without 2 μM Strep-His-*Xenopus* γTuNA. **See also Figure S1** – Addition of γTuNA to Xenopus egg extract does not affect γTuRC assembly or stability.

Structural studies of γTuRCs from yeast, frogs (*Xenopus laevis*), and humans revealed remarkable conservation of the γ-tubulin ring structure, although the composition of γTuRC differs substantially across these organisms (Kollman et al., 2015; Liu et al., 2020; Wieczorek et al., 2020; Consolati et al., 2020). Intriguingly, in all of these structures, the pitch and diameter of the γ-tubulin ring is incompatible with that of the assembled MT lattice. This suggests that γTuRC undergoes a conformational change to reduce its diameter before it can nucleate MTs (Thawani et al., 2020; Liu et al., 2020). One possibility is that this activating conformational change is stimulated by direct binding of “activation” factors. At the same time, other modes of activation are also plausible.

Here, we utilize *Xenopus* egg extract and *in vitro* single molecule imaging assays to demonstrate that the γTuNA domain directly interacts with γTuRC and identify the residues involved. Moreover, we find that γTuNA binds γTuRC as a dimer, triggering a sharp increase in γTuRC’s nucleation ability. This increase in γTuRC activity is sufficient to counteract indirect regulation by the tubulin-sequestering protein, stathmin. In sum, our study broadly reveals new insights into the spatiotemporal control of MT nucleation.

## 4. Results

### Cdk5rap2’s γTuNA domain increases MT nucleation in *Xenopus* egg extract

To study how Cdk5rap2 regulates γTuRC’s activity, we added Cdk5rap2’s purified γTuNA domain (Fig. 1A-B) to *Xenopus laevis* egg extract and assessed its impact on microtubule (MT) nucleation (Fig. 1C). Using total internal reflection (TIRF) microscopy and fluorescent end binding protein 1 (EB1) to label growing MT plus ends (Fong et al., 2016), we quantified individual MT nucleation events (Fig. 1C-D). In the control reaction, the egg extract showed a typical low level of MT nucleation (Fig. 1C, “buffer”, ~3 MTs per field).

In contrast, addition of wildtype γTuNA triggered an increase in MT nucleation of up to ~75-fold in a titration series (Fig. 1C-D). The γTuNA F75A mutant did not significantly increase MT number even at the highest concentration (3.6 *μ*M, Fig. 1C-D). Thus, γTuNA does indeed activate MT nucleation in extract and requires the F75 residue, confirming prior studies (Fong et al., 2008; Choi et al., 2010). Using sucrose gradients to fractionate mock and γTuNA-treated extracts, we also conclude that γTuNA has no effect on γTuRC assembly, ruling out one possible explanation for this increase in MT number (Fig. S1).

### γTuNA requires the universal MT template, the γ-tubulin ring complex (γTuRC)

We next asked whether γTuNA’s ability to increase MT nucleation in extract was dependent on the known MT nucleator, γTuRC. To do this, we depleted extracts of γTuRC via pulldown of γTuNA-coupled beads before repeating the TIRF assay. With a single round of depletion, we observed a loss of >75% of γ-tubulin signal indicating a depletion of γTuRC (Fig. 1E). In the mock-treated extract where γTuRC was not depleted, γTuNA’s ability to increase MT nucleation levels remained unchanged (Fig. 1F). In contrast, exogenous γTuNA no longer activated MT nucleation in γTuRC-depleted extracts (Fig. 1F). Hence, the γTuNA domain requires the universal MT template, γTuRC, to activate MT nucleation.

### The γTuNA domain can designate new artificial MTOCs by recruiting γTuRC

As γTuNA co-depletes γTuRC, we wondered whether this interaction would be sufficient to generate artificial MT asters (Fig. 2A). To that end, we coated micron-scale beads with wildtype or mutant γTuNA and added them to extract. After a pulldown step, we assayed these beads for MT aster formation *in vitro* in the presence of purified tubulin and GTP under oblique TIRF (Fig. 2B). We found that wildtype γTuNA-coated beads formed large MT asters mimicking the potent MT nucleation of the centrosome (Fig. 2B). In contrast, the F75A mutant beads formed severely impaired asters (Fig. 2B). Mock-treated beads did not form asters.

**Figure 2.**
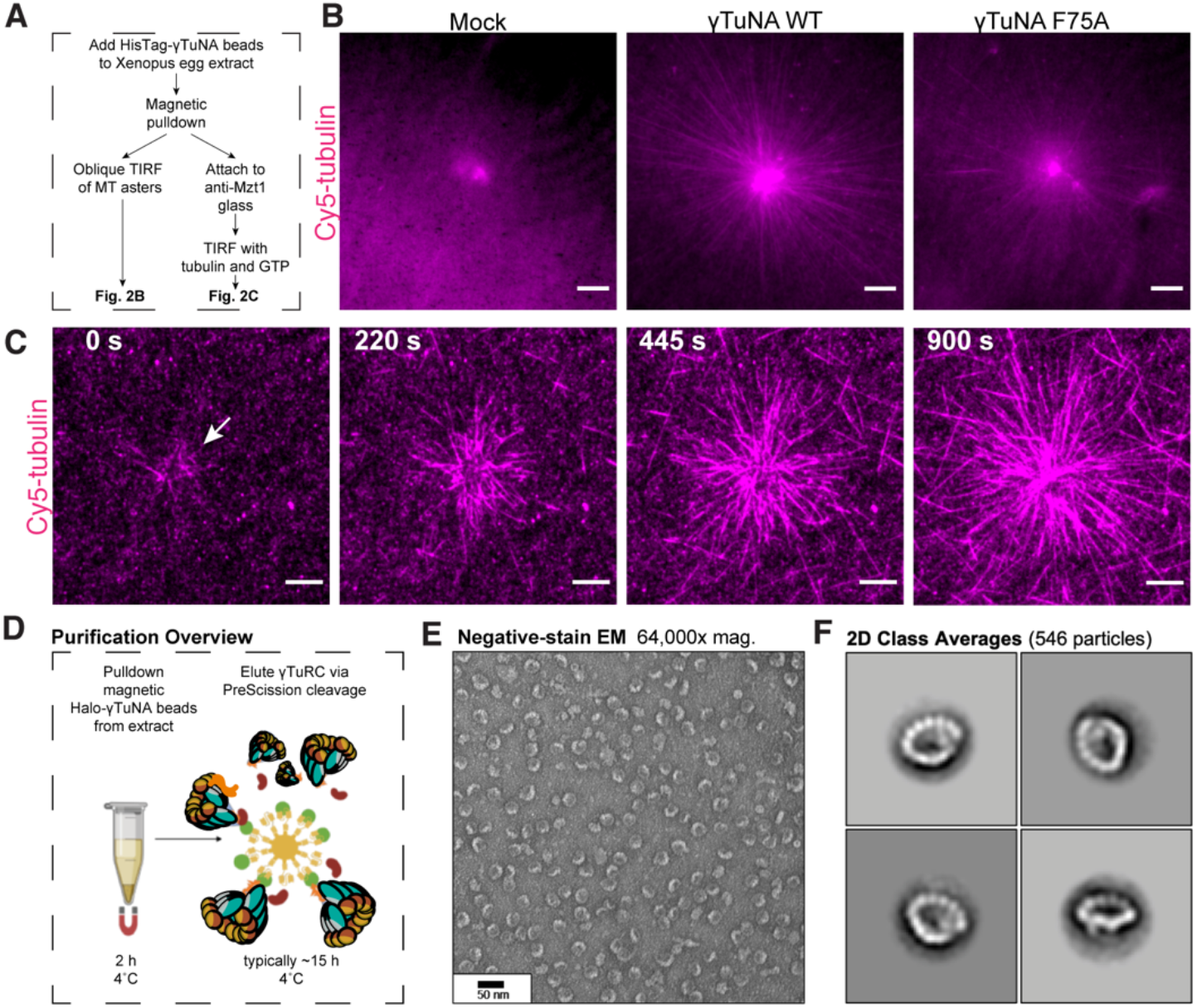
The γTuNA domain strongly recruits MT nucleation factors (including γTuRC) from Xenopus egg extract. **A**) Schematic of experiments for B and C. **B**) Oblique TIRF images of MT asters from beads *in vitro* after 10 min. HisPur magnetic beads coated with either bovine serum albumin (mock), Strep-His-*Xenopus* γTuNA wildtype (WT), or F75A mutant were incubated with extract, pulled-down, and washed. These were then diluted 1/1000 with polymerization mix containing 15 μM tubulin and 1 mM GTP, before imaging with TIRF. 5% Cy5-tubulin was used to label MTs. Bar = 5 μm. **C**) Time-lapse imaging of MT aster growth from wildtype γTuNA beads *in vitro*. As in part B, wildtype γTuNA beads were pulled-down from extract and washed. These were then incubated on DDS-surface treated coverslips containing anti-Mzt1 antibody to attach beads containing γTuRC. After a wash step, polymerization mix was added prior to time-lapse TIRF imaging. Frames are shown over the course of 15 minutes (900s). Bar = 5 μm. **D**) Diagram of major steps for purification of endogenous *Xenopus* γTuRC using magnetic beads coupled to Strep-His-HaloTag-3C-human γTuNA. **E**) Representative image of purified γTuRCs via negative-stain electron microscopy. Magnification is 64,000x, taken at 80 kV with a Philips CM100 transmission electron microscope. Bar = 50 nm. **F**) 2D class averages of 546 γTuRC particles picked from negative-stain EM images like in E. Each image represents one of four top classes. **See also Figure S2** – Mass Spectrometry (Quant-IP) reveals γTuRC is the dominant factor present after extract pulldown of γTuNA-beads. **See also Figure S3** – Purity and concentration assessment of γTuRCs purified via Halo-γTuNA. **See also Figure S4** – γTuRCs purified via Halo-γTuNA pulldown are fully assembled rings **Video-1** – Post-pulldown wildtype γTuNA beads nucleate asters in vitro.

To confirm the presence of γTuRC, we repeated the bead pulldown from extract and attached the beads via an antibody against Mzt1, a γTuRC subunit, to surface-treated coverslips (Fig. 2A). We then added fluorescent tubulin and GTP before live imaging via TIRF microscopy (Fig. 2C). Critically, we observed that wildtype γTuNA beads attached and formed large MT asters *in vitro*, indicating that these beads had retained γTuRC and any other necessary MT nucleation factors (Fig. 2C, Video-1). Thus, γTuNA, a relatively small peptide, is sufficient to specify new sites of γTuRC-mediated MT nucleation. Critically, this finding allowed us to develop a new γTuRC purification scheme based on scaled-up pulldowns with Halo-human γTuNA (outlined in Fig. 2D-F), which we discuss in more detail later.

### γTuNA is an obligate dimer

Having confirmed that γTuNA strongly recruits γTuRC from extract, we next investigated the γTuNA-γTuRC interaction. We selectively mutated hydrophobic residues found within a heptad-repeat region of γTuNA. In a recent structure of γTuRC, these residues were localized to a parallel coiled-coil structure that directly interacts with γTuRC (Wieczorek et al., 2020a). Specifically, we mutated the hydrophobic residues F63, I67, L70, and L77 to either alanine or aspartate (Fig. 3A). To compare the effect of these mutations at these well-conserved residues, we generated both human and *Xenopus laevis* versions of these constructs.

**Figure 3.**
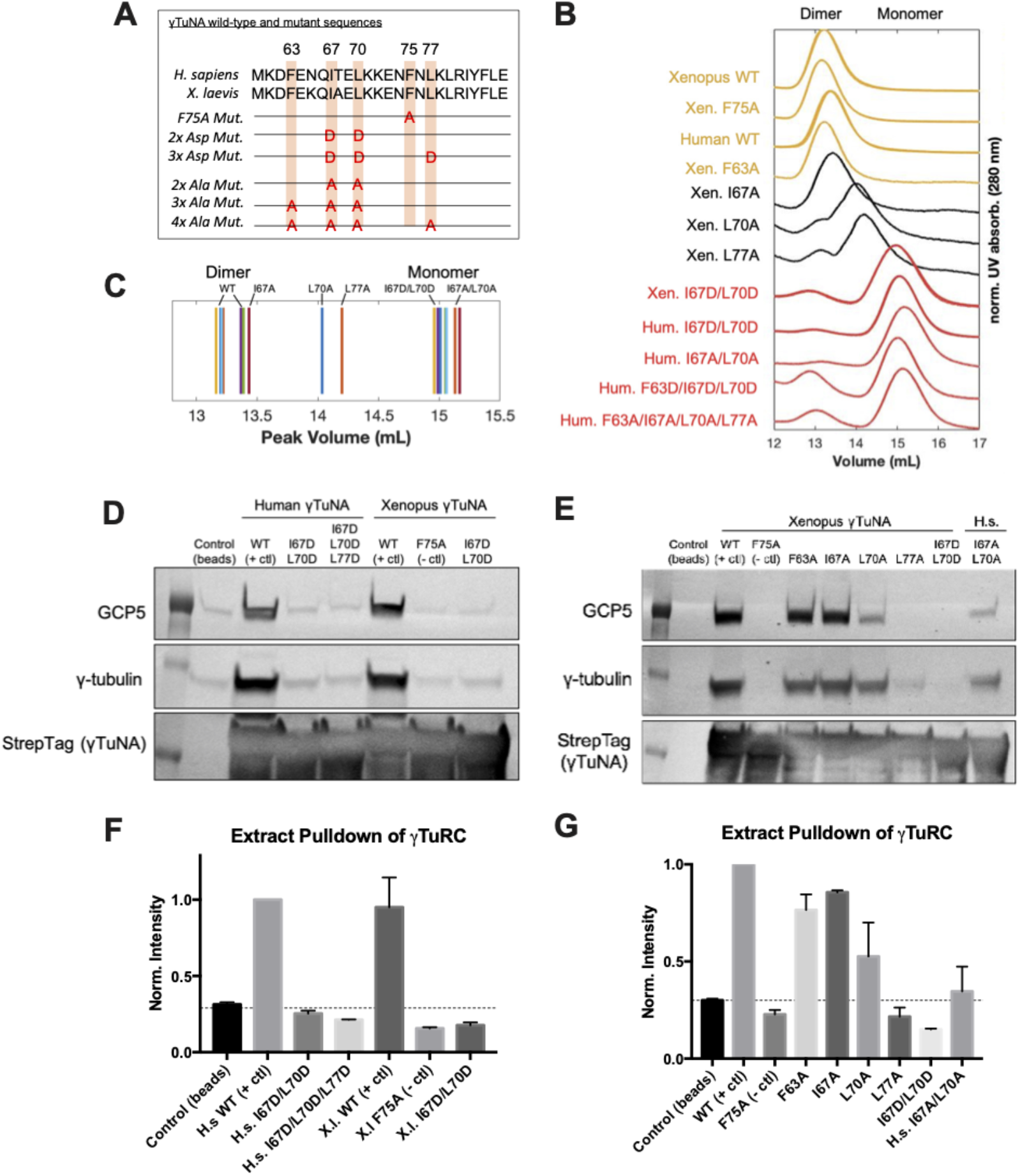
γTuNA requires both dimerization and the F75 residue to bind γTuRC in extract. **A**) Diagram of individual and combinatorial mutations made in human and *Xenopus* versions of Strep-His-Halo-3C-γTuNA (Halo-γTuNA) at residues F63, I67, L70, F75, and L77. **B**) Size-exclusion chromatography traces for human and *Xenopus* Halo-γTuNA wildtype and mutant constructs. Proteins were run at 50 μM on a Superdex 200 increase 10/300GL column (Cytiva) on an Äkta Pure system. Absorbance traces (A280 nm) were normalized by their peaks and plotted stacked as shown. **C**) Diagram of peak retention volumes for each construct tested. **D**) Western blots for γTuRC components, GCP5 and γ-tubulin, pulled down by beads coupled to human and *Xenopus* Halo-γTuNAs incubated in egg extract. The Strep-tag blot is shown as a bead loading control. **E**) Western blots as in D, except comparing pulldowns done with Halo-*Xenopus* γTuNA alanine point mutants, with wildtype and F75A mutants as positive and negative controls. **F**) Quantification of γTuRC pulldowns shown in D, normalized to the band intensity for human wildtype Halo-γTuNA beads. N =3. Error bars are s.d. **G**) Same quantification of γTuRC pulldowns as in F, except for pulldowns as done in E. Normalized to the band intensity of wildtype *Xenopus* γTuNA. N = 2. Error bars are s.d.

We initially focused on the double, triple, and quadruple mutants for both human and *Xenopus* γTuNAs. We performed size-exclusion chromatography (SEC) and compared the peak retention volumes of wildtype and mutated γTuNAs. Our SEC data revealed that wildtype γTuNA is a dimer (Fig. 3B-C). By comparing the SEC traces for the double, triple, or quadruple mutants from both *Xenopus* and human γTuNAs, we found that γTuNA dimerization was dependent on residues I67, L70, and L77 (Fig. 3B). The double hydrophilic mutants (I67D/L70D) from both human and *Xenopus* versions were entirely monomeric. This was also true for the human double-alanine mutant, I67A/L70A (Fig. 3B).

To resolve each residue’s individual contribution to γTuNA dimerization, we generated alanine point mutants for F63, I67, L70, and L77 in *Xenopus* γTuNA. We also tested the F75A mutant of *Xenopus* γTuNA, as we wanted to know whether its loss-of-function coincided with loss of dimerization. We compared the SEC traces for these point mutants and found that mutating residues F63 or F75 to alanine had no deleterious effect on γTuNA dimerization (Fig. 3B). In contrast, individually mutating residues I67, L70, or L77 increasingly interfered with dimerization, resulting in intermediate populations between full dimer and full monomer (Fig. 3B-C). Mutation of the L70 or L77 residues resulted in the most drastic impairment, further confirming that this central region is crucial for γTuNA dimerization.

### Both dimerization of γTuNA and its F75 residue are critical for binding γTuRC

With the insight that γTuNA is an obligate dimer, we next asked whether dimerization was required to bind γTuRC. We performed pulldowns of N-terminally Halo-tagged γTuNA mutants from *Xenopus* egg extract. We determined the amount of γTuRC bound for each γTuNA construct by probing for the γTuRC components GCP5 and γ-tubulin (Fig. 3D-G). We found that both human and *Xenopus* double aspartate mutants (I67D/L70D), as well as the human triple mutant (I67D/L70D/L77D) did not bind γTuRC, indicating that loss of dimerization results in loss of γTuRC binding (Fig. 3D,F). Interestingly, we found that the intermediate dimer mutants (I67A, L70A, or L77A) had correspondingly intermediate levels of γTuRC binding ability (Fig. 3E). The I67A mutant, for example, was weakly impaired in terms of dimerization (Fig. 3B) and subsequently retained its ability to bind γTuRC (Fig. 3E,G). As dimerization was increasingly impaired in the L70A and L77A mutants, γTuRC binding became increasingly weaker (Fig. 3G). In the most extreme example, the L77A mutant, which had the most substantial dimerization defect, had complete loss of γTuRC binding on par with the F75A negative control (Fig. 3G). Lastly, we also found that the human double alanine mutant (I67A/L70A) retained some ability to bind γTuRC. This residual amount of γTuRC signal was roughly the same as the single L70A mutant, again demonstrating the overwhelming contribution of the core L70 residue to dimerization, and now, to binding γTuRC (Fig. 3E,G).

### Rescuing dimerization of the γTuNA domain is not enough to rescue γTuRC binding

As the dimerization ability of the intermediate mutants correlated with their levels of γTuRC binding, we hypothesized that the degree of dimerization determined the amount of γTuRC binding. If true, then artificially rescuing dimerization should rescue full γTuRC binding. Furthermore in flies, the leucine zipper (LZ) domain in the Cdk5rap2 homolog, Centrosomin (Cnn), is known to dimerize (Feng et al., 2017; PReM domain in Fig 1A). When considering full-length Cdk5rap2, it is unclear if dimerization of γTuNA is enhanced by the downstream LZ dimerization domain. To understand the effect this LZ domain might have on γTuNA’s dimerization and γTuRC binding, we set out to rescue γTuNA dimerization by addition of a non-native LZ domain.

We fused γTuNA to the short LZ domain of the yeast transcription factor GCN4 (Schroeder et al., 2016), which forms a constitutive homodimer. After purification, we tested each construct via SEC to assess dimerization (Fig. 4). We found that all γTuNA-GCN4 fusion constructs formed dimers (Fig. 4A). We next performed pulldowns with these constructs from extract to assess whether forced dimerization now rescued γTuRC binding (Fig. 4B). Surprisingly, we found that rescuing dimerization did not rescue γTuRC binding (Fig. 4B), and the relative levels of γTuRC signal for each GCN4-fused mutant were essentially unchanged when compared to the mutants without GCN4 (Fig. 3E vs Fig. 4B). Thus, the ability of the γTuNA domain to dimerize does not require the presence of the downstream LZ domain, nor can the LZ compensate for deficiencies in γTuRC-binding caused by disrupting the γTuNA coiled-coil. Altogether, our SEC and pulldown data indicate that γTuNA is an obligate dimer that requires both efficient coiled-coil formation—depending mainly on L70 and L77—and F75 to bind γTuRC.

**Figure 4.**
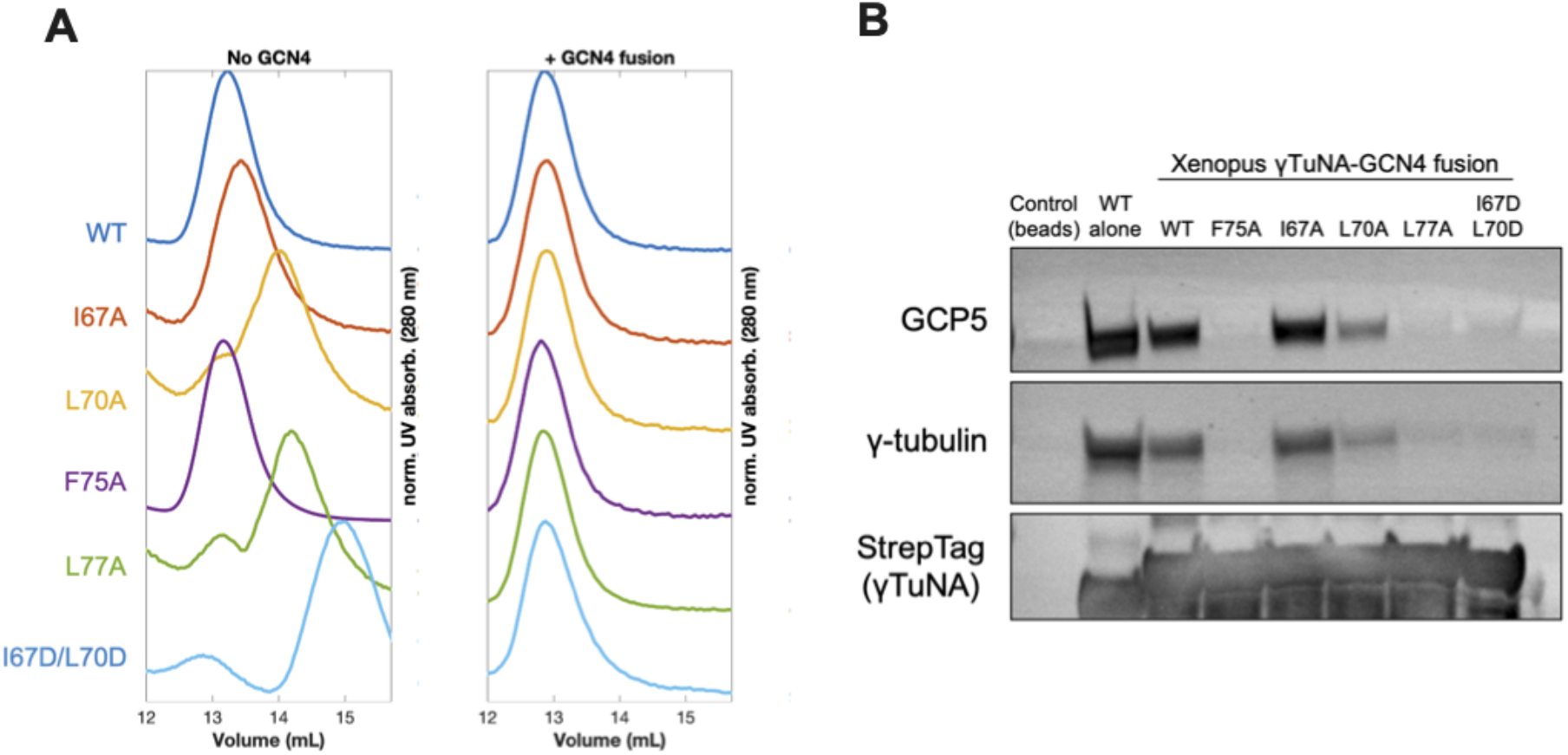
Rescuing γTuNA dimerization is not enough to rescue γTuRC binding. **A**) Size-exclusion chromatography traces for *Xenopus* Strep-His-Halo-3C-γTuNA (Halo-γTuNA) or Halo-γTuNA-GCN4 constructs. These constructs were either wildtype or single alanine mutations of residues I67, L70, F75, and L77. Traces were normalized by their maximum peak absorbance and plotted as shown. Proteins were run at 50 μM in a Superdex 200 increase 10/300GL column (Cytiva) on an Äkta Pure system. **B**) Western blot of γTuRC components, GCP5 and γ-tubulin, pulled down from extract by beads coupled to Halo-*Xenopus* γTuNA wildtype (as a positive control) or Halo-*Xenopus* γTuNA-GCN4 fusion constructs (wildtype, alanine point mutants, or an aspartate double mutant). StrepTag blot is shown as a bait loading control.

### Efficient γTuNA dimerization and the F75 residue are required for full γTuRC activation in extract

Having identified specific mutations that impaired γTuNA’s ability to dimerize and bind γTuRC, we next asked what effect these mutants had on MT nucleation in extract. We added wildtype or mutant *Xenopus* γTuNA to freshly prepared extracts and again tracked MT plus-ends via fluorescent EB1 as a measure of MT number (Fig. 5). As before, wildtype γTuNA triggered an increase in MT nucleation, when compared to the buffer control (Fig. 5). The F75A mutant had little effect on extract MT levels (Fig. 5). Similarly, the L77A mutant, which cannot bind γTuRC in extract (Fig. 3G), did not increase MT nucleation (Fig. 5). Intriguingly, however, when we examined the intermediate γTuRC-binding mutants I67A and L70A, we found that the I67A mutant activated MT nucleation to ~50% of wildtype levels, but L70A had no activity (Fig. 5B). Because I67A had only half the activity of wildtype despite retaining ~85% of the wildtype’s γTuRC-binding ability (Fig. 3G), γTuRC activation in extract has a non-linear relationship to γTuNA’s γTuRC binding. In addition, because the L70A mutant retained ~50% γTuRC binding ability (Fig. 3G) but had no activity in extract (Fig. 5B), it appears that there is a threshold to γTuRC’s activation in extract which only the wildtype and I67A γTuNA constructs can overcome.

**Figure 5.**
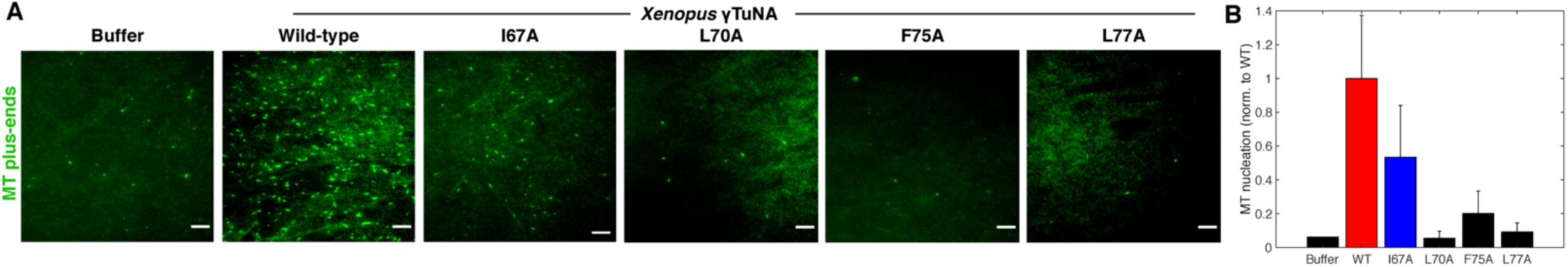
Complete γTuNA dimerization is required to maximally increase MT nucleation in extract. **A)** TIRF assay of MT nucleation in extract after addition of 2 μM wildtype or single alanine mutant versions of Strep-His-*Xenopus* γTuNA. EB1-mCherry was used to count MTs (MT nucleation) and is shown pseudo-colored green. Images were taken after 5 minutes at 18-20°C. Bar = 5 μm. **B**) Quantification of MT nucleation (MT number) normalized to the buffer condition, then normalized by the wildtype condition across four independent experiments. Red bar denotes wildtype level, while the blue bar denotes the effect of the I67A mutant. Error bars are s.d.

### The γTuNA domain directly activates MT nucleation by γTuRC *in vitro*

While we had explored the effect of wildtype γTuNA and its dimer mutants on MT nucleation in extract, we had yet to determine if γTuNA directly increased γTuRC’s activity *in vitro*. As we briefly mentioned (Fig. 2D-F), we used beads coupled to a Halo-human γTuNA construct to purify endogenous *Xenopus* γTuRC from extract (Fig. 2D-F), similar to previous work (Wieczorek et al., 2020b). Mass spectrometry confirmed that the dominant co-precipitant was indeed *Xenopus* γTuRC (Fig. S2). We also confirmed the presence of fully assembled γTuRC rings via negative-stain electron microscopy (Fig. 2E-F and Fig. S4).

Using this purified γTuRC, we investigated the effect of wildtype and mutant γTuNAs on γTuRC’s activity *in vitro* via single molecule imaging assays (Fig. 6). In these assays, biotinylated γTuRCs were attached to passivated coverslips before imaging with TIRF microscopy (schematized in Fig. S5). This not only offers high signal-to-noise but also allows tracking of individual γTuRC-mediated MT nucleation events.

**Figure 6.**
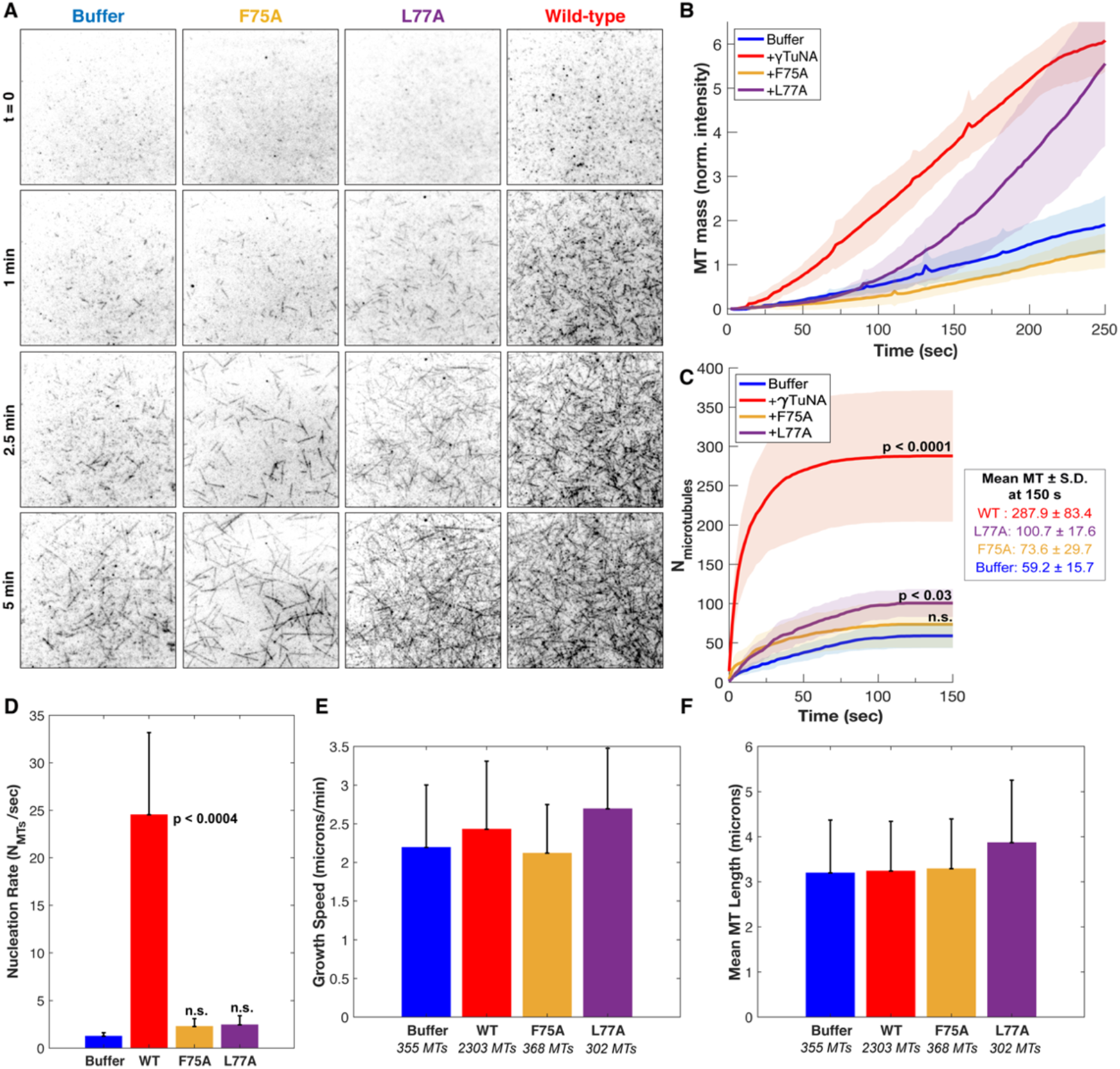
γTuNA dimers directly activate γTuRC MT nucleation ability in vitro. **A**) Single molecule TIRF assays of γTuRC-mediated MT nucleation *in vitro*. Purified *Xenopus* γTuRCs were biotinylated and attached to passivated coverslips via surface-bound Neutravidin molecules. Polymerization mix containing 15 μM tubulin,1 mM GTP, and either control buffer or 3.3 μM Strep-His-*Xenopus* γTuNA was then added. Wildtype, F75A, and L77A versions of γTuNA were tested. 5% Alexa 568-tubulin was used to visualize MTs. Images were taken every 2s, for 5 min total, at 33.5°C. Wildtype γTuNA (n = 8), buffer control (n = 6), γTuNA-F75A (n = 5), and γTuNA-L77A (n = 3). **B**) Quantification of total MT signal (MT mass) over time, normalized to the buffer condition at 150 s. **C**) Mean MT number over time (measured for the first 150 s). The box shows the mean MT number (± s.d.) at 150s for each condition. **B and C)** Solid lines are the mean over time, with error clouds representing standard deviation. **D**) Quantification of the initial nucleation rate (MTs nucleated per sec) for each condition. The curves used to calculate part C were fit to an exponential function to determine *k* (the nucleation rate); each *k* was then averaged; see methods. Buffer: 1.2 ± 0.37 MTs/sec, WT: 24.5 ± 8.6 MTs/sec, F75A: 2.3 ± 0.83 MTs/sec, L77A: 2.4 ± 0.96 MTs/sec. **C and D)** Two-sample unpaired t-tests were used to compare the buffer mean to the experimental means. P-values are noted if below p = 0.05. **E**) Mean MT growth speed for each condition. **F**) Mean maximum MT length for each condition. **E and F**) Bars are means, error bars are s.d. Wildtype γTuNA (n = 2303 MTs), buffer control (n = 355 MTs), γTuNA-F75A (n = 368 MTs), and γTuNA-L77A (n = 302 MTs). **See also Figure S5** – *Overview and additional single molecule TIRF data*. **See also Figure S6** – *Bulky N-terminal tags on γTuNA directly inhibit γTuRC activity in extract and in vitro*. **See also Figure S7** – *Simulation of γTuNA-dependent activation of γTuRC*. **Video-2**–Wildtype γTuNA directly stimulates γTuRC in vitro. **Video-3** – Video of simulated γTuNA-dependent activation of γTuRC.

We started by first comparing total MT mass generated in our assay (Fig. 6B). Strikingly, the addition of γTuNA triggered a ~5-fold increase in MT mass as compared to the buffer control (Fig. 6B, Video-2). To determine if this was a direct stimulation of γTuRC’s activity, we then quantified the number of γTuRC-nucleated MTs within the first 150 seconds (Fig. 6C), the MT nucleation rate (Fig. 6D), the mean MT growth speed (Fig. 6E), and the mean maximum MT length (Fig. 6F). These quantifications revealed that wildtype γTuNA sharply increased the γTuRC nucleation rate from 1.2 MTs/sec to 24.5 MTs/sec (~20-fold increase, Fig. 6C-D), with no significant effect on growth speed or MT length (Fig. 6E-F). We also found that wildtype γTuNA saturated our assay within 30 seconds (Fig. 6C), with a decreased nucleation rate of 0.15 MTs/sec that remained constant for the remainder of the experiment (Fig. S5). Thus, we conclude that γTuNA’s effect is almost exclusively due to a direct ~20-fold increase in γTuRC activity and not due to altered MT dynamics.

As expected, the F75A mutant did not increase MT mass *in vitro* (Fig. 6B). Furthermore, both the F75A and L77A mutants had no significant effect on the initial γTuRC nucleation rate, growth speed, or MT length (Fig. 6D-F). However, we did observe that L77A caused a weakly significant increase in MT number beginning at 150 seconds (Fig. 6C, p ~ 0.03). Similarly, we found that around 150 seconds L77A increased γTuRC’s nucleation rate 1.4-fold when compared to buffer (Fig. S5). Beyond 150 seconds, the L77A mutant triggered a delayed increase in MT mass (Fig. 6B), a behavior not observed in extract (Fig. 5). From this discrepancy, we inferred that the presence of other factors in extract blocks impaired dimer mutants like L77A from interacting with or stimulating γTuRC. We will return to L77A’s two-phase behavior below; regardless, our observations suggest that even in a purified (and interaction-permissive) setting, impaired dimerization prevents efficient γTuRC activation.

### Simulation of γTuNA-dependent activation of γTuRC *in vitro*

We were intrigued by L77A’s two-stage behavior and also wanted to confirm that wildtype γTuNA’s effect on MT mass could be modeled purely based on our measured γTuRC nucleation rates. To do this, we wrote a deterministic simulation for MT nucleation, with the only input being the initial nucleation rate (Fig. S7). To simplify our simulation, we assumed no MT catastrophes and no spontaneous nucleation. From this, we generated movies of simulated MT nucleation and growth (Fig. S7-A, Video-3). We also generated plots of simulated MT mass and MT number over time (Fig. S7-B,C).

We hypothesized that wildtype γTuNA activated γTuRC’s nucleation rate exclusively, and thus our simulated MT mass and MT number curves should match the curves in Fig. 6B and 6C. In our simulation, wildtype γTuNA generated a 6-fold increase in simulated MT mass, which closely matches our observed 5-fold increase (Fig. S7-B vs. Fig. 6B).

Our simulation also predicted the MT mass and MT number curves for the F75A and buffer conditions (Fig. S7-B-C vs Fig. 6B-C). However, because our simulation is determined solely by the initial nucleation rates, it failed to predict the late stage behavior of the L77A mutant, and simulated only a 1.8-fold increase in MT mass, compared to our observed 3 to 5-fold increase (Fig. S7-B vs Fig. 6B). Instead, altering our simulation such that L77A switched at 150 seconds from its initial nucleation rate to the wildtype rate better captured the 3-fold increase in MT mass (Fig. S7D-E vs. Fig. 6B). We hypothesize that this could be thought of as a switch from a previously impaired dimer state to a complete dimer (L77A vs wildtype, Fig. 3B).

Altogether, we conclude that complete dimerization of γTuNA is required to activate γTuRC in extract. Our *in vitro* data suggests this requirement for efficient dimerization is due to competition or regulation by other factors in extract, as even the impaired dimer mutant L77A can activate γTuRC in a minimal, purified system. We propose that the two phases of L77A mutant activity could arise from sequential binding of two L77A monomers, where γTuRC activation occurs only once the second monomer binds (schematized in Fig. S7F). Critically, from both our *in vitro* data and simulations, we conclude that wildtype γTuNA directly activates γTuRC, stimulating its nucleation ability ~20-fold without affecting MT dynamics.

### γTuNA activation of γTuRC in extract overcomes the effect of negative regulators like stathmin

As there appeared to be an activation barrier in extract, but not *in vitro*, we investigated whether known negative regulators of MT nucleation were responsible. We focused on the tubulin-sequestering protein stathmin (or op18), which regulates the available tubulin pool for nucleation and polymerization (Belmont and Mitchison, 1996). For every mole of stathmin present (1.5 *μ*M endogenous concentration), two moles of tubulin are removed (Gigant et al., 2000; Wühr et al., 2014).

We sought to first confirm that stathmin negatively regulated MT nucleation by γTuRC, and then determine whether γTuNA had any effect on this. To that end, we added increasing amounts of exogenous stathmin to extract and measured its effect on MT nucleation (Fig. 7A-B). At double and triple the endogenous concentration of stathmin in extract (~3 *μ*M or ~4.2 *μ*M final), we observed a complete loss of MT nucleation and polymerization (Fig. 7A-B). Surprisingly, γTuNA was still able to activate MT nucleation (Fig. 7A), even at the highest concentration of stathmin tested, with a ~17-fold increase in the number of MTs (Fig. 7B, 2.7 *μ*M stathmin).

**Figure 7.**
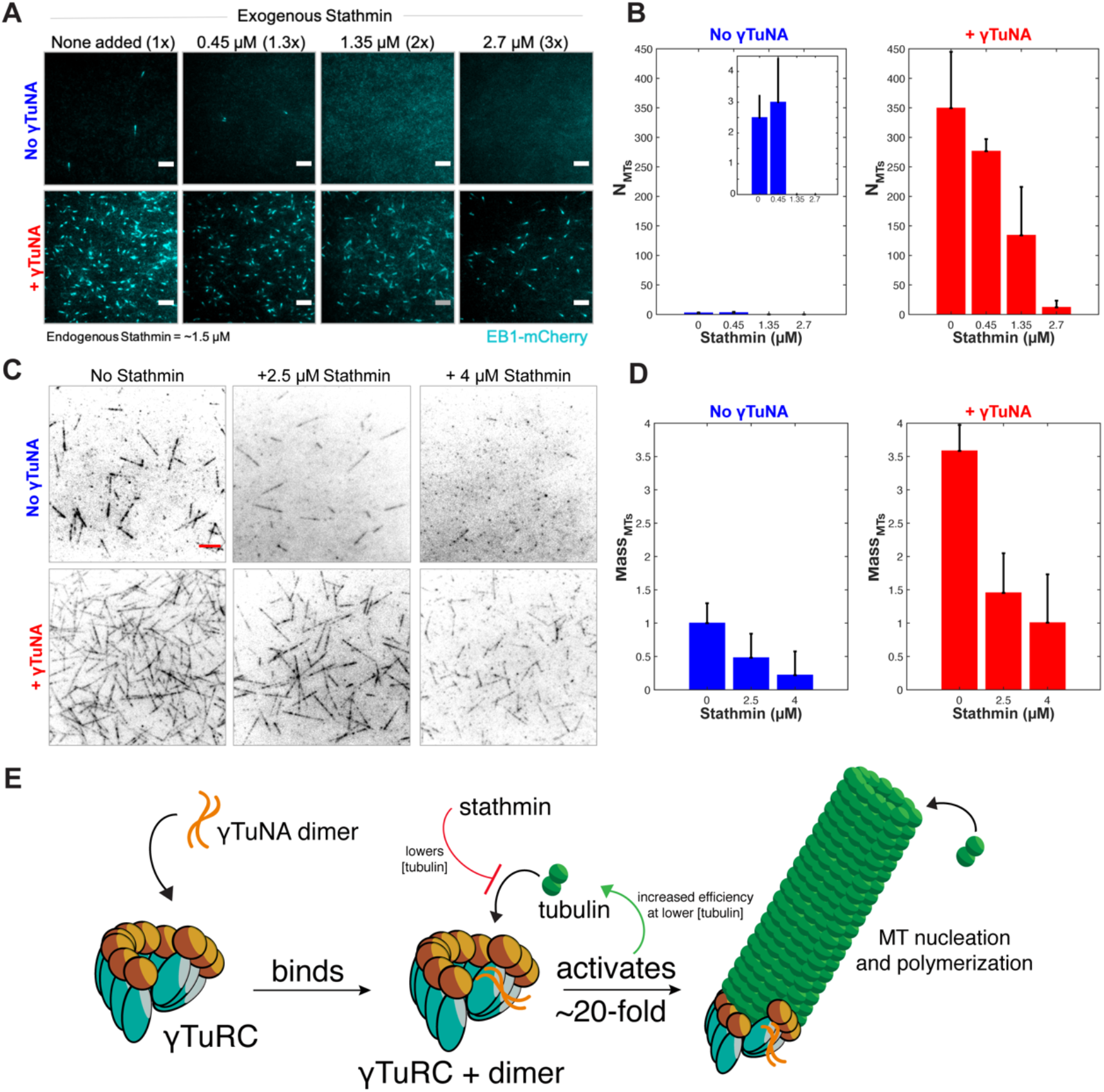
γTuNA activation of γTuRC overcomes stathmin repression in extract and in vitro. **A**) TIRF assay of extract MT nucleation (Strep-His-γTuNA vs stathmin). His-SNAP-tag-*Xenopus* stathmin (isoform 1A) was added at 0.45, 1.35, and 2.7 μM final concentration to extract. Either buffer control or 2.3 μM Strep-His-*Xenopus* wildtype γTuNA was also added. EB1-mCherry was added to visualize MT plus-ends (MT number). Images were taken after 5 min at 18-20°C and pseudo-colored cyan. Bars = 5 μm. **B**) Quantification of MT number for each concentration of stathmin tested, n = 3; bars are s.d. **C**) Single molecule TIRF assay of γTuRC MT nucleation *in vitro*. Purified γTuRCs were assayed at 33.5°C with 15 μM tubulin, 1 mM GTP, either alone, in the presence of 2.5 μM or 4 μM stathmin, or with 3.3 μM Strep-His-*Xenopus* γTuNA. 5% Alexa-568-tubuin was used to visualize MTs. Red bar = 5 μm. **D)** Quantification of MT mass present at 250 seconds (intensity normalized to the buffer control) from C. Bars are mean normalized intensity; error bars are s.d. N = 3 for all conditions, except: 4 μM stathmin alone, n = 4; and, 4 μM stathmin with γTuNA, n = 5. **E)** Summary model of γTuNA-mediated activation of γTuRC’s MT nucleation ability.

Next, we assessed whether the γTuNA-γTuRC complex could also overcome stathmin’s effect *in vitro* (Fig. 7C). For simplicity, we tracked the total MT signal (or MT mass) produced after 250 seconds in the TIRF-based nucleation assay (Fig. 7D). We first observed the activity of γTuRC alone at 15 *μ*M tubulin (Fig. 7C, upper left panel). We next tested γTuRC in the presence of either 2.5 or 4 *μ*M stathmin and found that stathmin resulted in losses of 52% and 78% MT mass, respectively (Fig. 7C, “No γTuNA” conditions). Interestingly, addition of γTuNA to stathmin and γTuRC rescued MT mass (Fig. 7C-D, “+ γTuNA”). In fact, at 4 *μ*M stathmin with γTuNA, MT mass levels were restored to the level of the γTuRC control (no stathmin, no γTuNA).

Furthermore, as it is known that stathmin is phosphorylated to release tubulin (Gavet et al., 1998, Kuntziger et al., 2001), we asked whether γTuNA increased the levels of phosphorylated stathmin. However, Western blots of extract treated with γTuNA showed there was no appreciable difference in stathmin phosphorylation (data not shown). Altogether then, our findings suggest that γTuNA counteracts the effect of stathmin in extract by increasing the efficiency of γTuRC-mediated MT nucleation at lower tubulin concentrations.

## 5. Discussion

### A model for γTuNA-mediated activation of γTuRC

In this work, we addressed how MT nucleation is regulated in space and time by investigating Cdk5rap2’s γTuNA-mediated activation of γTuRC (Fig. 7E). We showed that γTuNA is an obligate dimer and that dimerization is crucial for binding γTuRC, which is consistent with a recent structure of γTuRC bound to a γTuNA dimer (Wieczorek et al., 2020a). Moreover, we defined the core residues required for both γTuNA dimerization and subsequent γTuRC binding. We found that γTuNA dimers directly activate γTuRC-dependent MT nucleation in extract and *in vitro*. Finally, we uncovered that γTuNA overcomes barriers to MT nucleation posed by the tubulin sequestrator stathmin. Because γTuNA domains are also found in myomegalin and the branching factor TPX2, among others, our findings are broadly applicable to multiple MTOCs and model eukaryotes from yeast to humans.

It remains an open question how the γTuNA dimer directly enhances γTuRC’s nucleation activity. It is tempting to speculate that γTuNA binding triggers a conformational change of γTuRC from its wide diameter lattice (Liu et al., 2020; Wieczorek et al., 2020b; Consolati et al., 2020) to a closed state. Because previous work has characterized the structures of γTuNA bound to human and *Xenopus* γTuRC and observed no obvious structural change (Liu et al. 2020, Wieczorek et al. 2020b), it is possible that binding a γTuNA dimer only transiently biases γTuRC towards the closed ring conformation. Prior modeling suggested that the free energy provided by stochastic binding and lateral association of the first 3-4 tubulin dimers is sufficient to overcome the energy barrier between γTuRC’s open and closed states (Thawani et al., *eLife*, 2020). Activation factors such as γTuNA may be able to reduce the energy barrier required for ring closure. In parallel, locally elevated tubulin concentrations, combined with a subtle bias toward γTuRC ring closure, might be sufficient to trigger nucleation. In support of this, several nucleation sites, such as the centrosome and TPX2-based branch sites, locally concentrate tubulin creating an environment in which γTuNA’s effect could be further enhanced (Baumgart et al., 2019; M. King et al., 2020). Ultimately, detection of a potential low-population activated state of γTuRC may require high levels of tubulin and GTP combined with sophisticated structural methods.

Altogether, our model for γTuNA activation of γTuRC involves a direct binding event between a γTuNA dimer and γTuRC, which activates γTuRC’s MT nucleation ability ~20-fold (Fig. 7E). At the same time, γTuNA-containing proteins like Cdk5rap2 bind specific MTOCs (like the centrosome), thus recruiting poised γTuRCs to areas of high tubulin concentration. Furthermore, these newly localized γTuNA-γTuRC modules can overcome regulation by stathmin. This sets the stage for subsequent MT nucleation and helps ensure γTuRC’s activity is spatiotemporally restricted.

### Resolving conflicting data concerning γTuNA’s effect on γTuRC

This study was partly motivated by an apparent discrepancy between the original reports of γTuNA’s ability to activate γTuRC *in vivo* and *in vitro* (Fong et al., 2008; Choi et al., 2010) and more recent *in vitro* data from our group and others that found little to no effect (Liu et al., 2020; Thawani, Rale et al., 2020).

In their recent structural study of antibody-purified *Xenopus* γTuRC, Liu and colleagues concluded that the N-terminal region of Cdk5rap2 (or CEP215) containing the γTuNA domain had little to no effect on γTuRC MT nucleation *in vitro* (Extended Data Fig. 9b from Liu et al., 2020). They did, however, report that wildtype γTuNA (CEP215N) co-precipitated γ-tubulin, while the F75A mutant did not (Extended Data Fig. 9c from Liu et al., 2020). Liu and colleagues used N-terminally GST tagged versions of γTuNA (CEP215N). Like our colleagues, we find that the presence of a large N-terminal tag does not interfere with γTuNA’s ability to bind γTuRC (Fig. 3). However, our studies revealed that a bulky N-terminal Halo-tag on γTuNA turns this activator into a specific inhibitor of γTuRC-mediated MT nucleation in extract and *in vitro* (Fig. S6). This is likely due to the steric clash produced by two copies of the bulky N-terminal tag in proximity to the critical nucleation interface on the γ-tubulin ring. We note that the original reports from the Qi group used the small FLAG tag (Choi et al., 2010), and our work is based on the small Strep-His tag at the N-terminus.

In our prior work, we had established an antibody-based *Xenopus* γTuRC purification, which was pioneering at that time, yet with limited yield and batch-to-batch variability. Although N-terminally 6xHis-tagged γTuNA activates MT nucleation in extract (as presented in Fig. 1 of this study), it had little to no effect on the original antibody-purified *Xenopus* γTuRC *in vitro* (Fig. 6 in Thawani, Rale et al., 2020). Since then, we have developed a purification method based on human Halo-γTuNA, which is routinely at least 20-fold higher yield, higher purity, and ultimately has more robust activity. This resulted in increased density of nucleation competent γTuRCs present in our single molecule assays, as well as better detection of γTuNA’s activation effect. As such, we validate and extend the original γTuNA studies by the Qi group.

### The role of cis-regulatory elements in γTuNA-containing proteins

It is known that human CDK5RAP2’s γTuNA/CM1 domain (51-200 aa; 149 aa total) is phosphorylated, with evidence suggesting this phosphorylation is required for binding and activation of γTuRC in cells (Hanafusa et al., 2015). Our version of γTuNA (46 aa total) does not contain the known S140 phospho-target, yet still binds and stimulates γTuRC in extract and *in vitro*. This suggests that the larger 51-200 aa version of γTuNA contains an auto-inhibitory sequence that is relieved by phosphorylation. In support of this, a recent report found that the *C. elegans* homolog of Cdk5rap2, SPD-5, is phosphorylated downstream of its γTuNA domain to allow binding with γTuRC (Ohta et al., 2021). The authors speculated that the N-terminal γTuNA domain is folded over and blocked from interacting with other binding partners, until phosphorylation by PLK1 relieves this auto-inhibition (Ohta et al., 2021). We suggest that, if correct, this proposed mechanism might also prevent γTuNA dimerization by blocking efficient coiled-coil formation between monomers. Thus, these cis-regulatory elements might help prevent ectopic activation of γTuRC.

### Other factors possibly involved in tuning γTuNA-γTuRC activity

Our mass spectrometry analysis revealed that γTuRC is the dominant co-precipitant for wildtype versions of human and *Xenopus* γTuNA (Fig. S2). We also detected the nucleoside diphosphate kinase 7 (better known as NME7), which validates prior work reporting its presence as either a γTuRC binding factor or possibly as a key activating subunit (Liu et al., 2014). Surprisingly, we detected three unique proteins that were enriched at a higher level than NME7 (Fig. S2-F) and had not been reported to directly interact with γTuRC or γTuNA. These were the cyclin-dependent kinase 1 (CDK1) subunits A and B, as well as the type II delta chain of the calmodulin-dependent protein kinase (CAMK2D). Hence, these might be novel co-factors for γTuRC.

While our work has now revealed that γTuNA-containing proteins can directly activate the MT nucleation template, γTuRC, several questions remain. Chief among these is whether the co-nucleation factor, XMAP215/ch-TOG, which is now known to act synergistically with γTuRC to nucleate MTs (Thawani, Kadzik, et al., 2018), might further enhance γTuNA’s effect on γTuRC. Or in a similar vein, what effect does the aforementioned γTuRC subunit NME7 have on this process? Investigating how multiple factors simultaneously tune γTuRC activity is an exciting avenue that can further our understanding of microtubule nucleation.

## Supporting information

MATLAB-Numerical Simulation of gTuNA-gTuRC MT nucleation

MATLAB- Graphical Simulation of gTuNA-gTuRC MT nucleation

## 6. Acknowledgments

The authors would like to thank all members of the Petry lab, past and present. In particular, we would like to thank Dr. Raymundo Alfaro-Aco. We would also like to thank Bernardo Gouveia for his insightful comments and Dr. Jodi Kraus for sharing reagents. We especially thank both the Princeton Mass Spectrometry core facility and the ThermoFisher Center for Multiplexed Proteomics (TCMP) at Harvard Medical School. We also thank the Imaging and Analysis Center (IAC) at Princeton University, which is partially supported by the Princeton Center for Complex Materials (PCCM), a National Science Foundation (NSF) Materials Research Science and Engineering Center (MRSEC; DMR-2011750), and in particular, Dr. John Schreiber. We also thank Dr. Ron Vale (UCSF) for the plasmid containing GCN4 (Addgene plasmid# 74608).

## Funding

MR was supported by a Howard Hughes Medical Institute (HHMI) Gilliam fellowship and a National Science Foundation (NSF) Graduate Research fellowship. This work was also supported by an NIH New Innovator Award 1DP2GM123493, the Pew Scholars Program in the Biomedical Sciences (00027340), and the David and Lucile Packard Foundation 2014–40376 (all to SP).

## 7. Author contributions

Conceptualization, M.R. and S.P.; Methodology, M.R. and S.P.; Investigation, M.R., B.R., B.P.M, S.M.T., and S.P.; Writing – Original Draft, M.R. and S.P.; Writing – Review & Editing, M.R., B.R., B.P.M, S.M.T. and, S.P.; Visualization: M.R. and S.P.; Funding Acquisition, M.R. and S.P.; Supervision, S.P.

## 8. Declaration of interests

The authors declare no competing interests.

## 9. Materials and methods

### 9.1. Cloning and Purification of human and *Xenopus* γTuNA constructs

The fragment of human CDK5RAP2 (51-100) containing the CM1 motif/γTuNA domain was sub-cloned into a bacterial expression pST50 vector using Gibson cloning. This vector was engineered with N-terminal Strep-TagII, 6xHis, TEV cleavage, HaloTag, and PreScission 3C protease cleavage sites. This vector was then truncated to remove all but the first 46 N-terminal codons (see Table 1).The resulting construct, Strep-His-TEV-HaloTag-3C-human γTuNA, (Halo-human γTuNA), was expressed in Rosetta 2 (DE3) *E. coli* cells. Rosetta 2 cells were grown in 2 L terrific broth (TB) cultures to O.D. = 0.7 and induced with 0.5 mM IPTG at 16°C for 18h. The cultures were pelleted, snap-frozen, and stored at −80°C.

**Table 1.**
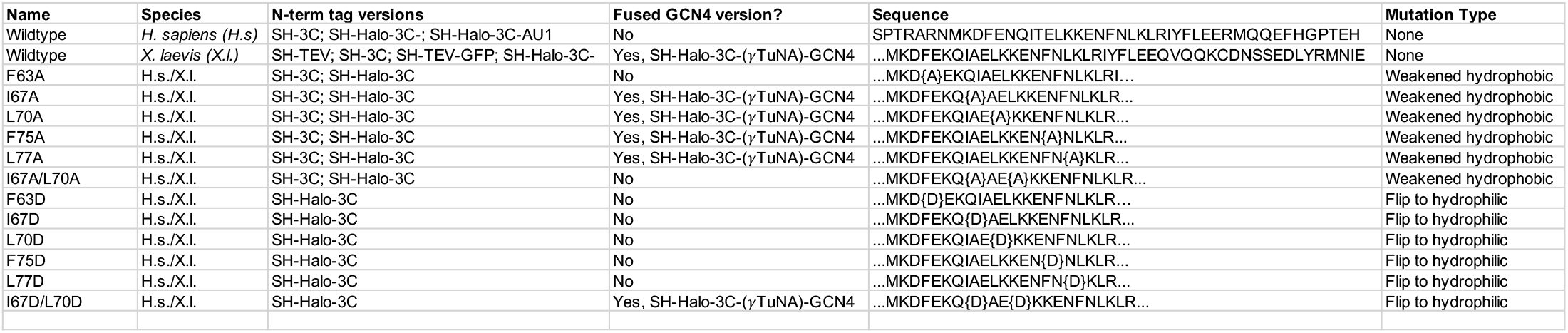
γTuNA constructs generated in this study. Key: S = StrepTagII, H = 6xHis-tag, 3C = human rhinovirus 3C (PreScission) protease cleavage site, TEV = tobacco etch virus protease cleavage site, GCN4 = yeast transcription factor GCN4 dimerization domain. Mutated residues in the “Sequence” column are designated using { } brackets.

The CM1/γTuNA-motif in *Xenopus laevis* Cdk5rap2 was confirmed via sequence alignment to the human version and inserted using Gibson cloning into the same pST50 bacterial construct as above. This generated Strep-His-TEV-HaloTag-3C-*Xenopus* γTuNA (Halo-Xenopus γTuNA). The loss-of-function mutation, F75A, first identified by Fong et al., 2008, was introduced into the Halo-Xenopus γTuNA sequence at the equivalent, conserved phenylalanine to make Strep-His-TEV-HaloTag-3C-*Xenopus* γTuNA F75A (Halo-Xenopus γTuNA F75A). Both these constructs were expressed as described above with the human version.

The amino acid sequence of the human γTuNA used is: SPTRARNMKDFENQITELKKENFNLKLRIYFLEERMQQEFHGPTEH. The sequence for the *Xenopus* γTuNA (wildtype) used in this work is: MKDFEKQIAELKKENFNLKLRIYFLEEQVQQKCDNSSEDLYRMNIE.

All γTuNA constructs generated in this study are listed in Table 1. For GCN4 C-terminal fusions, we used pET28a-Hook3 aa 1-160-GCN4 plasmid, which was a gift from Dr. Ron Vale (Addgene plasmid # 74608).

To purify the human and *Xenopus* Halo-γTuNA constructs, 2 L TB cell pellets were thawed on ice and resuspended into 50 mL of Strep Lysis Buffer (50 mM TRIS, pH = 7.47, 300 mM NaCl, 6 mM *β*-mercaptoethanol, 200 *μ*M PMSF, 10 *μ*g/mL DNase I, and a single dissolved cOmplete EDTA-free Protease Inhibitor Cocktail tablet (cat # 11873580001, Roche). Cells were resuspended using a Biospec Tissue Tearor (Dremel, Racine, WI) and lysed in an Emulsiflex C3 (Avestin, Ottawa, Canada) by processing four times at 10,000-15,000 psi. Cell lysate was spun at 30,000 rpm for 30 min, 2°C in a Beckman Optima-XE 100 ultracentrifuge, 45Ti rotor. Supernatant was then passed twice through a 15 mL column volume (CV) of Strep-Tactin Superflow resin (IBA, Goettingen, Germany). The column was then washed with 10 CV of Strep Bind buffer (50 mM TRIS, pH = 7.47, 300 mM NaCl, 6 mM *β*-mercaptoethanol (BME), 200 *μ*M PMSF). The γTuNA proteins were then eluted with 1.5 CV of Strep Elution buffer (Strep Bind Buffer with 3.3 mM D-desthiobiotin (cat. #2-1000-005, IBA)). Yield and purity were assessed via SDS-PAGE gel and Coomassie stain. Concentration was assessed via Bradford assay. All γTuNA constructs yielded between 40 and 60 mg of protein (per 2 L TB culture) at >98% purity.

### 9.2. Size-exclusion chromatography of Halo-γTuNA proteins

For all size-exclusion assays, we used an ÄKTA Pure system with a Superdex 200 increase 10/300 GL column (cat. #28-9909-44, Cytiva, Marlborough, MA), with a 500 *μ*L manual injection loop. All assays were done in CSFxB, 6 mM BME, without sucrose at 4°C. Strep-His-TEV-Halo-3C-γTuNA constructs shown in Fig. 3 were run at ~50 *μ*M final concentration in a total volume of 550 *μ*L Strep bind buffer (see above), at a 0.7 mL/min flow rate. Absorbance at 280 nm was used to track the protein peak. Each trace was normalized by the maximum peak for that run, prior to combined plotting with all other constructs in MATLAB (ver. R2019a, MathWorks, Natick, MA).

### 9.3. TIRF imaging of MT nucleation in *Xenopus laevis* egg extracts

*Xenopus laevis* egg extracts were prepared as previously described (Good and Heald, 2018). For assaying γTuNA’s effect on MT nucleation levels, 7.5 *μ*L of extract were incubated on ice with 0.5 *μ*L 10 mM Vanadate (0.5 mM final), 0.5 *μ*L 1mg/mL end-binding 1 (EB1)-mCherry protein (0.05 mg/mL final), 0.5 *μ*L of 1 mg/mL Cy5-tubulin (0.05 mg/mL final), 0.5 *μ*L of CSFxB (10% sucrose), and 0.5 *μ*L of TRIS control buffer (50 mM TRIS, pH = 7.47, 300 mM NaCl, 6 mM *β*-mercaptoethanol) or 0.5 *μ*L of γTuNA protein (previously diluted in Tris control buffer such that final concentrations are as stated in Figure 1). Reactions (10 *μ*L total) were gently mixed by pipetting once, before adding to a channel on a 6-channel slide at 18-20°C. All γTuNA concentrations for each condition (wildtype or F75A mutant) were imaged in parallel on the same slide. Multi-channel images were acquired sequentially and at 1 min intervals using the NIS-Elements AR program (NIKON, ver. 5.02.01-Build 1270). The 647 nm/Cy5 channel (excitation: 678 nm, emission: 694 nm) was used for microtubules (MT) and the 561 nm channel for EB1 MT plus-tips (ex: 587 nm, em: 610 nm). The images were captured on a Nikon Ti-E inverted system, with an Apo TIRF 100x oil objective (NA = 1.49), and an Andor Neo Zyla (VSC-04209) camera with no binning and 100 msec exposures. Resulting images were 2048 by 2048 pixels (132.48 *μ*m x 132.48 *μ*m). The 561 nm channel was pseudo-colored green.

To quantify MT nucleation levels, we extracted the number of MT plus ends (tracked by EB1 spots) for each condition using the 561 nm/EB1-mCherry channel. We wrote a macro in FIJI (ImageJ; Schindelin et al., *Nature Methods*, 2012) to automate counting EB1 spots. Briefly, the 561 nm channels were contrast matched (same minimum and maximum values for LUTs) for all conditions run on the same day, with the same extract. Next, 50 *μ*m by 50 *μ*m (800 x 800 pixels^2^) representative windows were cropped from each field of view, smoothed using the FIJI function (“Process—> Smooth”), and thresholded using the Yen option. The built-in FIJI functions “Find edges” and “Analyze particles” were then used to count the number of thresholded spots. These values, representing the number of MT plus ends, were then normalized to the buffer control for each condition to obtain the fold change in MT number. These fold changes were then averaged across four trials and plotted using Prism GraphPad 7 (GraphPad Software, San Diego, CA). Representative images are shown in Fig. 1.

#### Stathmin Assays in Extract

For assaying γTuNA dimer mutants’ effects on MT nucleation, we used the above procedure except all constructs shown in Fig. 5 were tested at 2 *μ*M final concentration using 600 msec exposures of EB1 (561 nm channel) only. For assaying γTuNA’s effect on stathmin in extract (Fig. 7), we used the same procedure above except we added either 0.5 *μ*L unlabeled 6xHisTag-SNAP-Stathmin or Tris control buffer instead of 0.5 *μ*L CSFxB. Data were plotted in MATLAB.

### 9.4. TIRF imaging of MT nucleation from γTuNA-coated beads *in vitro*

#### γTuNA Bead Assay (endpoint version)

We first saturated 5 *μ*L of micron-scale, HisPur Ni-NTA magnetic beads (cat. # 88831, ThermoFisher) with 50 *μ*L of either bovine serum albumin (6.5 mg/mL BSA, as mock), wildtype *Xenopus* Strep-His-γTuNA (71 *μ*M), or *Xenopus* Strep-His-γTuNA F75A mutant (~117 *μ*M) in CSFxB buffer. After 30 min incubation on ice, beads were removed with a magnet, washed with 150 *μ*L CSFxB, and resuspended with 50 *μ*L BRB80 buffer (80 mM PIPES, pH = 6.8 with KOH, 1 mM MgCl_2_, 1 mM EGTA). These beads were then diluted 1/1000 in polymerization mix (15 *μ*M total tubulin with 5% labeled Cy5-tubulin and 1 mM GTP in BRB80 buffer) and added to a channel on a glass slide. Beads were located via differential interference contrast (DIC) microscopy. Then MT aster formation for each condition was imaged via oblique TIRF microscopy at 5 min intervals up to 25 minutes.

#### Anti-Mzt1 γTuNA Bead Assay (live imaging version)

We first passivated glass coverslips with dichlorodimethylsilane (DDS, cat. #440272-100ML, Sigma), as previously published (Gell et al., 2010, Alfaro-Aco et al., 2017). These coverslips were then attached with double-sided tape to glass slides to create multi-channel imaging chambers. To each chamber, we added in order: 1) 50 *μ*L of BRB80, 2) 20 *μ*L of 375 *μ*g/mL Mzt1 antibody in BRB80 (anti-Mzt1, cat. # ab178359, Abcam), 3) 1% Pluronic-127 (cat. # P6866, ThermoFisher) in BRB80, and 4) 10 *μ*L of 1/1000 diluted wildtype γTuNA beads in BRB80 (pulled from extract). At each step, we paused for 5 min incubations at room temperature. Just prior to imaging via TIRF, we then added 10 *μ*L of ice-cold BRB80, followed by cold polymerization mix (15 *μ*M total tubulin with 5% labeled Cy5-tubulin and 1 mM GTP in BRB80). Images were taken at room-temperature (18-20°C).

### 9.5. Purification of native *Xenopus* γ-tubulin ring complex (γTuRC) via Halo-human γTuNA pulldown

To purify native *Xenopus* γTuRC from *Xenopus* egg extract, we employed a strategy similar to previous work (Wieczorek et al., *Cell*, 2020) and originally observed by Choi et al., *JCB*, 2010. Here we similarly use γTuNA as a bait for γTuRC, except we use the human version of Halo-γTuNA directly coupled to beads via the affinity of the Halo-tag for its substrate. These beads are then used to pulldown γTuRC from *Xenopus laevis* egg extract.

*Xenopus laevis* egg extracts were prepared as described previously. Extracts were snap-frozen in liquid nitrogen and stored at −80°C. One day prior to purification, 15 mg of Halo-human γTuNA were thawed and diluted to ~1 mg/mL in a 15 mL conical tube with Coupling Buffer (20 mM HEPES, pH = 7.5 with KOH, 75 mM NaCl) to a final NaCl concentration of 100-135 mM NaCl. Next, 2 mL of Halo Magne bead slurry (cat. #G7287, Promega, Madison, WI) were washed with MilliQ water and 3 CV of modified CSF-XB, 2% sucrose buffer (10 mM HEPES, pH = 7.7, 100 mM KCl, 1 mM MgCl2, 0.1 mM CaCl2, 5 mM EGTA, 2% sucrose). Washed beads were then incubated under constant rotation with the 15 mL of Halo-human γTuNA for 16 hours overnight at 4°C. The beads were collected next day via a magnetic stand and washed with 3 CV of CSF-XB and resuspended to 2 mL with CSF-XB, 2% sucrose. Beads were either used fresh or stored for up to a week at 4°C.

The next day, frozen extract (4 mL) was thawed in a room-temperature water bath. 500 *μ*L of clean, uncoupled Halo Magne beads were washed with MilliQ water and 3 CV of CSF-XB, 2% sucrose for pre-clearing of extract. Thawed extract was then incubated with the 500 *μ*L of uncoupled Halo beads for 30 min at 4°C with rotation to remove non-specific binders. The pre-clearing beads were removed with a magnet, and the now “pre-cleared” extract was used to resuspend the 2 mL equivalent of coupled Halo-human γTuNA beads. Halo-human γTuNA beads were incubated with the extract for 2 h, 4°C, while rotating. The beads were then removed from the extract, washed with 4.5 CV of CSF-XB, and if required, incubated with 40 *μ*M NHS-PEG4-Biotin (cat. #A39259, ThermoFisher) for 1h on ice and then resuspended with 2 mL of 3C Elution buffer (600 *μ*g PreScission protease diluted in CSF-XB, 2% sucrose). Proteins were eluted from the beads overnight at 4°C, 15-18h, with rotation.

The elution containing cleaved γTuNA, γTuRC, and other factors, was concentrated to ~300 *μ*L in a 100 kDa MWCO Amicon 4 mL spin concentrator. This was then spun through a 10%-50% w/w sucrose gradient (in CSFxB buffer) using a TLS55 rotor in a Beckman Coulter Optima MAX-XP ultracentrifuge at 200,000g, 2 °C for 3h. The gradient was manually fractionated such that the first fraction was the same size as the input (~300 *μ*L), and each subsequent fraction was 140 *μ*L in size. Samples of each fraction were run on a 4-12% Bis-TRIS SDS-PAGE gel for 10 min, 140V. The SNAP i.d. 2.0 rapid Western blotting system (cat. #SNAP2MM, EMD-Millipore, Burlington, MA) was used to determine the peak γTuRC fraction (usually Fraction 7; γ-tubulin, GTU-88 mouse antibody, cat. # T6557, Sigma, St. Louis, MO). A 5 *μ*L sample of the peak fraction was then added to glow discharged copper CF400-CU EM grids (cat. # 71150, Electron Microscopy Sciences, Hatfield, PA) for 1 min, and stained with 0.75% Uranyl Formate (UF) for 40 seconds. The grid was then imaged on a Philips CM100 transmission electron microscope at 64,000x magnification, 80 kV, to verify the presence of intact γTuRCs, as well as to assess purity. A representative image is shown in Figure 2. The peak γTuRC fraction was stored on ice and used within 24 h or snap-frozen and stored at −80°C. Our prep yielded an average peak concentration between 150-200 nM γTuRC. Via a fused Halo-AU1 reporter version of human γTuNA, we measured 50 nM γTuNA dimer in the peak γTuRC fraction, suggesting 66% of purified γTuRCs lost their γTuNA dimer bait at the end of the sucrose gradient step (Fig. S3).

### 9.6. Negative stain EM data processing

Data processing of negative-stain EM data was done using Relion 3.1 (Zivanov et al., 2018). We manually picked 4,866 total particles from 59 micrographs, followed by particle extraction and 5 rounds of 2D class averaging for particle sorting prior to 3D reconstructions. 4,593 particles were used for *ab initio*, non-templated reconstructions. Particles were further sorted with 3D class averaging, and 2,692 selected particles were used for structure refinement. Final refinement iterations were done using a 100 Å low-pass filtered mask with a contour cutoff of 0.007. This gave a final mask-sharpened map at 28 Å resolution as determined by the gold-standard FSC cutoff of 0.143 (Fig. S4). CryoEM movie data was aligned and CTF corrected using CryoSparc 3.2 (Punjani et al., 2017), followed by subsequent manual picking of particles on CTF-corrected micrographs. A total of 800 particles were manually picked, extracted, and used to calculate 2D class averages.

### 9.7. Pull-downs of γTuRC from *Xenopus* egg extract via Halo-γTuNA constructs

To compare γTuRC binding across mutant versions of γTuNA, we performed pulldowns from *Xenopus* egg extract with Halo-Magne beads (cat # G7281, Promega) coupled to N-terminally Halo-tagged versions of either human wildtype, *Xenopus* wildtype, or *Xenopus* dimer mutant γTuNA (as in Fig. 3). Mock beads (blocked with bovine serum albumin) were used to assess background levels of non-specific precipitants. For each condition, we diluted 2 mg total of bait protein in 1 mL of Coupling Buffer (see section 9.5). Next, we resuspended 170 *μ*L worth of Halo-Magne beads (washed 3x with Coupling Buffer) with each protein mix. These were incubated under rotation at 4°C for 1h. Beads were then collected via a magnetic stand, washed three times with Coupling Buffer, and then resuspended to 1 mL final volume with CSFxB (2% sucrose).

Next, 2 mL total of frozen *Xenopus* egg extract were thawed in a room-temperature water bath. Beads for each condition were then collected via a magnetic stand and resuspended with 200 *μ*L of extract. To each, we then added 800 *μ*L of CSFxB (2% sucrose), and incubated under rotation at 4°C for 2h. Beads were then washed twice with 1 mL of CSFxB (2% sucrose), prior to resuspension in 120 *μ*L of 1x SDS-PAGE sample buffer (6 mM DTT). Beads were then boiled at 95°C for 5 min. After magnetic removal of beads, the elutions were spun at 17,000g for 1 minute to remove aggregates or beads.

We ran 40 *μ*L of each elution per lane in 4-12% Bis-Tris SDS-PAGE gels at 140V for 1h. Proteins were then transferred to nitrocellulose membranes and probed for γTuRC components via Western blot: GCP5 (1:250 dilution of mouse anti-GCP5 antibody (E-1), # sc-365837, Santa Cruz Biotechnology, Santa Cruz, CA) and γ-tubulin (1:1000 dilution of GTU-88 mouse antibody, cat. # T6557, Sigma). To confirm equal coupling of bait to beads, we also probed for the Strep-tag on γTuNA (1:1000 dilution of *Strep-tag* mouse monoclonal antibody, cat. # 34850, Qiagen). Band intensities were measured in ImageJ and normalized in Prism 7 by the wildtype γTuNA band (positive control). Independent experiments, after normalization, were averaged together into the charts seen in Fig. 3 (N =3 for Fig. 3F, N=2 for Fig. 3G).

### 9.8. Single molecule TIRF imaging of γTuRC MT nucleation *in vitro*

#### Preparation of functionalized coverslips and imaging chambers

We utilized a previously published method to generate functionalized glass coverslips for single molecule TIRF imaging (Thawani et al., 2020; Consolati et al., 2020). Briefly, after sonication with 3 M NaOH, coverslips were sonicated with Piranha solution (2:3 ratio of 30% w/w H_2_O_2_ to sulfuric acid) to remove all organic residues. After washes with MilliQ water, we dried and then treated the coverslips with 3-glycidyloxypropyl trimethoxysilane (GOPTS, cat. # 440167, Sigma) at 75°C for 30 min. Unreacted GOPTS was removed with two sequential acetone washes, and the coverslips were then dried with nitrogen gas. Between sandwiched coverslips, we melted a powder mix of 9 parts HO-PEG-NH_2_ and 1 part biotin-CONH-PEG-NH_2_ by weight (cat. #103000-20 and #133000-25-20, Rapp Polymere, Tübingen, Germany) at 75°C. We pressed out air bubbles and repeated cycles of 75°C incubation and pressing until sandwiches were clear. After an overnight incubation at 75°C, the sandwiches were separated and washed in MilliQ water. These were then spun dry in air and stored at 4°C for up to 1 month.

We made imaging chambers by first making a channel on a glass slide with double-sided tape. To this channel, we added 2 mg/mL PLL-g-PEG (SuSOS AG, Dübendorf, Switzerland) in MilliQ water. After a 20 min incubation, the channel was rinsed thoroughly with MilliQ water and dried with nitrogen gas. Using a diamond pen, functionalized coverslips were cut into quarters. Each quarter piece was then added to the double sided tape on the slide with the functionalized surface facing down. Chambers were made fresh on the day of each experiment.

#### Attaching biotinylated, Halo-prepped γTuRC to functionalized chambers

As previously described (Thawani et al., *eLife*, 2020), imaging chambers were blocked first with 50 *μ*L of 5% w/v Pluronic F-127, then 100 *μ*L of assay buffer (BRB80 with 30 mM KCl, 1 mM GTP, 0.075% w/v methylcellulose 4000 cp, 1% w/v D-glucose, 0.02% w/v Brij-35, and 5 mM BME). Next, we added 100 *μ*L of casein buffer (0.05 mg/ml κ-casein in assay buffer). Here, we modified our protocol from our previous study by adding 20 *μ*L of 0.05 mg/mL NeutrAvidin (A2666, ThermoFisher), not 50 *μ*L of 0.5 mg/mL. We also decreased the cold block incubation at this step from 3 min to 1.5 min. We then washed the channel with 100 *μ*L of BRB80. Similarly, we diluted peak fraction Halo-prepped, biotin-γTuRC between 1/100 to 1/300 in BRB80 depending on the prep yield, not 1/5 as previously published. This was due to the increased yield of Halo-purified γTuRC as compared to our previous antibody-based method. We added 20 *μ*L of this γTuRC dilution and incubated for 7 min at room temperature. Finally, the chamber was washed with 20 *μ*L of cold BRB80. All buffers referenced here, except Pluronic F-127 (room-temperature), were used at 2°C.

#### Microtubule nucleation assay with purified γTuRC

Concurrently, we prepared our nucleation mix by mixing 15 *μ*M total unlabeled bovine tubulin (PurSolutions, Nashville, TN) with 5% Alexa 568-tubulin, 1 mg/mL BSA (cat. # A7906, Sigma) in assay buffer on ice. To this mix, we also added either Tris control buffer or Strep-His-3C-*Xenopus* γTuNA constructs (to 3.3 *μ*M final concentration). This was then centrifuged in a TLA100 rotor (Beckman Coulter) for 12 min at 80,000 rpm to remove aggregates. Finally, we added 0.68 mg/ml glucose oxidase (cat. # SE22778, SERVA GmbH, Heidelberg, Germany), 0.16 mg/ml catalase (cat. # SRE0041, Sigma). This nucleation mix was then added to the chamber with attached γTuRC and imaged immediately. For assays using stathmin/op18, we adjusted our nucleation mix so that it contained either stathmin control buffer (50 mM Tris, pH = 7, 300 mM NaCl, 400 mM imidazole) or 2.5 *μ*M or 4 *μ*M final concentration of 6xHis-SNAP-*Xenopus* stathmin (isoform 1A).

We used the same imaging set-up as our previous work (Thawani et al., 2020), notably a Nikon Ti-E inverted stand with an Apo TIRF 100x oil objective (NA = 1.49) and an Andor iXon DU-897 EM-CCD camera with EM gain set to 300. We again used an objective heater collar (model 150819–13, Bioptechs) to maintain 33.5°C for our experiments. However, for this study we captured time-lapse movies of the tubulin 561 nm channel at 1 frame every 2s for 5 min. All movies start within 1 min of the addition of ice-cold nucleation mix to the imaging chamber. Repeats were done with independent Halo-γTuRC preps: wildtype γTuNA reactions (n = 8), buffer control (n = 6), γTuNA-F75A (n = 5), and γTuNA-L77A (n = 3).

### 9.9. Analysis of single molecule TIRF MT nucleation assays

#### Analysis of total MT mass

For total MT mass measurements, we wrote a FIJI/ImageJ macro to measure the total 561 nm signal intensity for each frame in our time-lapses. To do this, the macro first filtered each frame using the Otsu method in the “Adjust Threshold” function to remove most background signal. Next, it used the “Measure” function and recorded the integrated density measurements for each frame. In MATLAB, we then background subtracted and normalized the integrated density value for each condition by the buffer control. We then plotted total MT mass over time, as shown in Fig. 6B. For the assays in Fig. 7C-D, we used the same method, except we plotted the total MT mass generated at 250 seconds, normalized by the buffer condition.

#### Analysis of MT number over time, nucleation rate, growth speed, and mean MT length

In each field of view, an area 40 *μ*m x 40 *μ*m (252 x 252 pixels^2^) was analyzed for the first 150 seconds of each reaction. We also corrected for minor translational drift in our movies by using the StackReg plugin for ImageJ (Thévenaz et al., 1998). Next, we wrote two FIJI/ImageJ macros to semi-automate our data analysis. The first macro generated kymographs (space-time plots) for each individual MT in the time-lapse, although each MT was manually selected. With the second macro, we manually extracted relevant parameters from these kymographs.

First, if the MT was spontaneously nucleated, the resulting kymograph would display bi-directional growth over time (appearing like a scalene triangle). If the MT was nucleated by a γTuRC, then one end did not grow over time resulting in kymographs with only a single growing edge (right angle triangle). Using the macro, we recorded whether each MT was spontaneously or γTuRC nucleated. If the MT was γTuRC-nucleated, we proceeded with our measurements. We next manually recorded the nucleation point (or origin) for each MT. We then drew a line along the growing edge and extracted its slope to generate the growth speed for that MT. We also measured the MT’s maximum length. These measurements were then imported into MATLAB (R2019a) and averaged across all reactions for each condition. The mean and standard deviation (s.d.) for the number of MTs nucleated over time were plotted using MATLAB, as shown in Fig. 6C. To determine the γTuRC nucleation rate, we fit all MT nucleation curves for each condition to Equation 1,

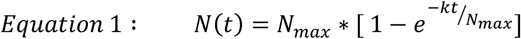

where *N(t)* = the number of MTs nucleated at that time point, *N_max_* = the maximum number of MTs nucleated after 150 sec, and *k* = the γTuRC nucleation rate. We used the nonlinear least squares fitting algorithm from MATLAB’s “lsqcurvefit” function to determine both *N_max_* and *k*. This nucleation rate (*k*) was then averaged for all reactions in each condition, including calculating the standard deviation of *k*. Both the mean and s.d. for the γTuRC nucleation rate are shown in Fig. 6D. We also plotted the linear slope of the curves at saturation (from 25 to 150 s) in Fig. S5. The mean and s.d. for the growth speed and mean maximum MT length are shown in Fig. 6D-E. These plots are not normalized and display the raw mean ± s.d. for each condition. Two-sample unpaired t-tests were used to determine if the means for each condition were significantly different from the buffer condition.

### 9.10. Simulations of γTuNA’s effect on γTuRC MT nucleation

To simulate the effect of γTuNA in our single molecule TIRF assay, we wrote a deterministic simulation in MATLAB based on the measured nucleation rates for each condition. For simplicity, we assume no spontaneous MT nucleation and no MT catastrophes. This system was modeled by Equation 2, as follows:

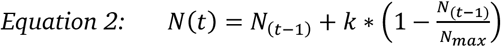

where *N(t)* = the current number of nucleated MTs (or active γTuRCs), *k* = the nucleation rate, and *N_max_* is the total number of γTuRCs activatable by that condition. At *N*(*t* = *0*), the number of active γTuRCs or nucleated MTs is zero. At each time step (1 sec), new MTs are added to the system according to the nucleation rate measured experimentally (*k*), which decreases until reaching saturation. These new MTs were then randomly placed on a simulated 40 x 40 pixel^2^ plane with a random initial orientation based on one of eight discrete conditions. At each new time step, the length of previous MTs is incremented by the average growth speed (across all conditions, in *μ*m/sec). This process of nucleation and growth is repeated until the end of the simulation, generating simulated movies of this process. For the L77A mutant, we also generated a second two-step simulation where, after 150 seconds, *k_L77A_* is arbitrarily redefined as the wildtype *k* (*k_WT_*), and *N_max_* in Equation 2 is redefined as (wildtype *N_max_* – L77A N(150s)), where L77A N_(150s)_ = the number of MTs already present at 150 seconds. Plots and simulated frames are shown in Fig. S7A.

### 9.11. Mass spectrometry identification of unique *Xenopus laevis* γTuNA binding factors

#### Sample Preparation

We performed pulldowns from *Xenopus* egg extract with Halo-Magne beads (Promega cat # G7281, Madison, Wisconsin, USA) coupled to either human wildtype, *Xenopus* wildtype, or *Xenopus* F75A mutant versions of Halo-γTuNA (Fig. S2). Uncoupled beads were used to assess background levels of non-specific precipitants. After washing, bound proteins were eluted with Glutathione-S-transferase (GST) tagged PreScission (HRV 3C) protease. Elutions were then subjected to a reverse GST step to remove most GST-PreScission. Ten percent of each elution sample was run on an SDS-PAGE gel and stained with Coomassie to confirm low levels of non-specific protein binders (bead control, Fig. S2-B). We also probed these elutions for the presence of γTuRC components, GCP5 and γ-tubulin, confirming they were only present when the bait was a wildtype version of γTuNA (Fig. S2-C). This pulldown was performed two times independently to generate a set of six samples (two replicates for each γTuNA condition) that were then submitted to the ThermoFisher Center for Multiplexed Proteomics (TCMP) for multiplexed quantitative mass spectrometry (Harvard University Medical School, Cambridge, MA).

#### Quantitative Mass Spectrometry (MS; Quant-IP)

At TCMP, the concentrations of our six samples were measured via a Pierce micro-BCA assay. Samples were then reduced with DTT and alkylated with iodoacetimide. This was followed by a protein precipitation step using methanol/chloroform. The resulting pellets were resuspended in 200 mM EPPS, pH 8.0. Samples were then digested sequentially using LysC (1:50) and Trypsin (1:100), determined by the protease to protein ratio. The digested peptides from each condition were then separately labeled with one of six tandem mass tags (TMT) for multiplexing (TMT-126, TMT-127a, TMT-127b, TMT-128a, TMT-128b, TMT-129a). All samples were then combined and run through basic pH reverse phase (bRP) sample fractionation utilizing an 8-to-28% linear gradient of acetonitrile (ACN; in 50 mM ammonium bicarbonate buffer, pH 8.0). These fractionated, TMT-tagged peptides were then analyzed via three sequential mass spectrometry scans (LC-MS^3^): a precursor ion Orbitrap scan (MS1), followed by an ion trap peptide sequencing scan (MS2), and a final Orbitrap scan to quantify the reporter ions (MS3).

#### Database Search Parameters

All MS2 spectra were analyzed using the Sequest program (Thermo Fisher Scientific, San Jose, CA, USA). Sequest was used with the following search parameters: peptide mass tolerance = 20 ppm, fragment ion tolerance = 1, Max Internal Cleavage Sites = 2, and Max differential/Sites= 4. Oxidation of methionine was specified in Sequest as a variable modification. MS2 spectra were searched using the SEQUEST algorithm with a Uniprot composite database derived from the *Xenopus* proteome containing its reversed complement and known contaminants. Peptide spectral matches were filtered to a 1% false discovery rate (FDR) using the target-decoy strategy combined with linear discriminant analysis. Identified proteins were filtered to a < 1% FDR. Proteins were quantified only from peptides with a summed SN threshold of >100 and MS2 isolation specificity of 0.5. From this, 21,902 unique peptides were detected, resulting in 3,214 total proteins. After filtering, this resulted in 2,842 unique, quantified proteins across our six γTuNA samples. The top 12 proteins (with at least 5 unique peptides) specifically enriched in the wildtype γTuNA samples are presented in Figure S2-F.

## Supplemental Figures

**Figure S1.**
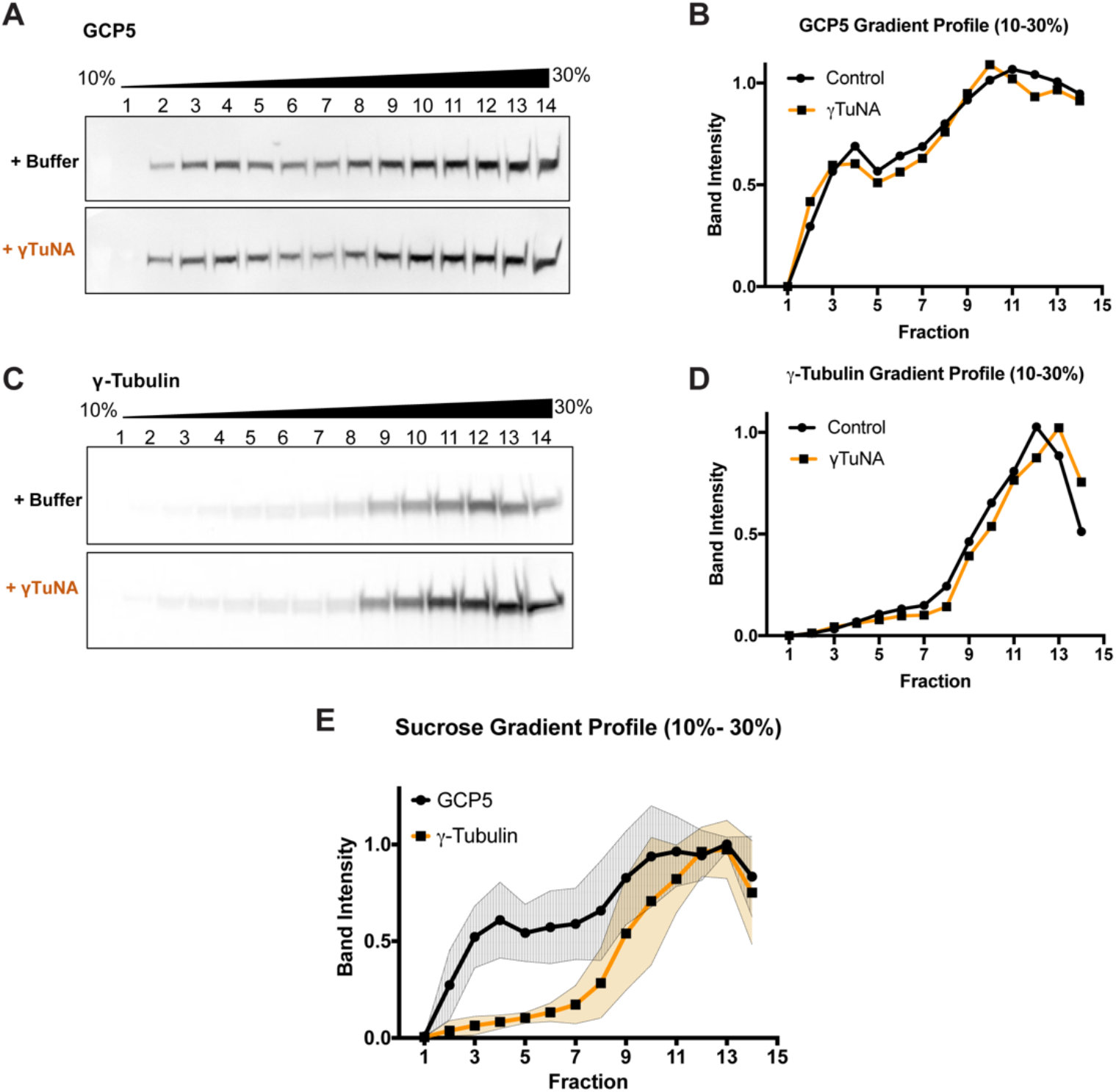
Addition of γTuNA to Xenopus egg extract does not affect γTuRC assembly or stability. High-speed centrifuged *Xenopus* egg extracts were run on 10-30% w/w sucrose gradients and fractionated. Western blots of GCP5 and γ-tubulin were quantified to generate a sucrose gradient profile for each target, as shown in A. GCP5 appeared to be in two separate populations: uncomplexed around fraction 3, and another in the γTuRC complex (around fraction 10-11). **A and C**) Western blots of GCP5 and γ-tubulin in gradient fractionated extracts treated with either buffer or 6 μM Strep-His-*Xenopus* γTuNA for 1h on ice. The presence of γTuNA did not shift intensity from the soluble peak of GCP5 (fraction 3) to the γTuRC fraction (fraction 11), nor did it have any effect on the distribution of γ-tubulin. **B and D**) Normalized band intensity profiles for GCP5 and γ-tubulin. **E**) Mean gradient profile for GCP5 and γ-tubulin with standard deviation (N = 2).

**Figure S2.**
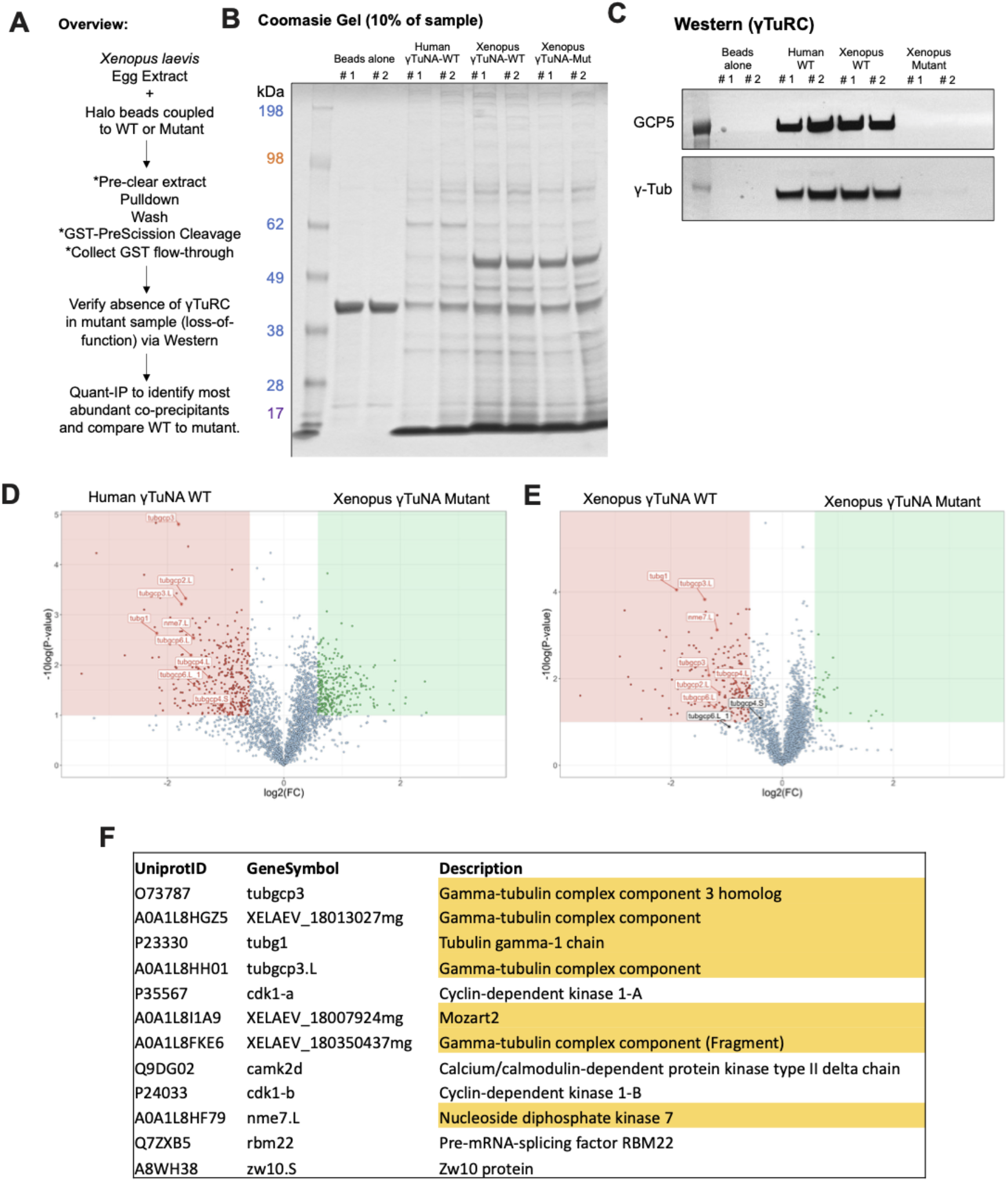
Mass Spectrometry (Quant-IP) reveals γTuRC is the dominant factor present after extract pulldown of Halo-γTuNA beads. **A**) Experiment overview of extract pulldowns with beads coupled to mock, human wildtype, *Xenopus* wildtype, and *Xenopus* F75A mutant versions of gTuNA. Bound factors were eluted using PreScission cleaveage of Halo-3C-γTuNA bait. **B)** SDS-PAGE and Coomassie stain of 10% of elution sample. The band at ~40 kDa corresponds to GST-PreScission. The band at ~7 kDa (dominant band < 17 kDa) corresponds to cleaved γTuNA bait. The dominant band at ~50 kDa was identified as *Xenopus* formimidoyltransferase cyclodeaminase (FTCD). **C**) Western blot of γTuRC components, GCP5 and γ-tubulin, in the elution samples. Note that only the wildtype γTuNA baits had detectable γTuRC signal. **D and E**) Volcano plots of proteins specifically enriched in the human or *Xenopus* γTuNA baits, as compared to the *Xenopus* F75A γTuNA mutant bait. **F**) Table displays the top 12 proteins specifically enriched by wildtype γTuNA bait, after subtraction of proteins present in F75A mutant bait. If a protein was not present in both human and Xenopus wildtype conditions or if it was identified with fewer than 5 unique peptides, it was not included in our analysis.

**Figure S3.**
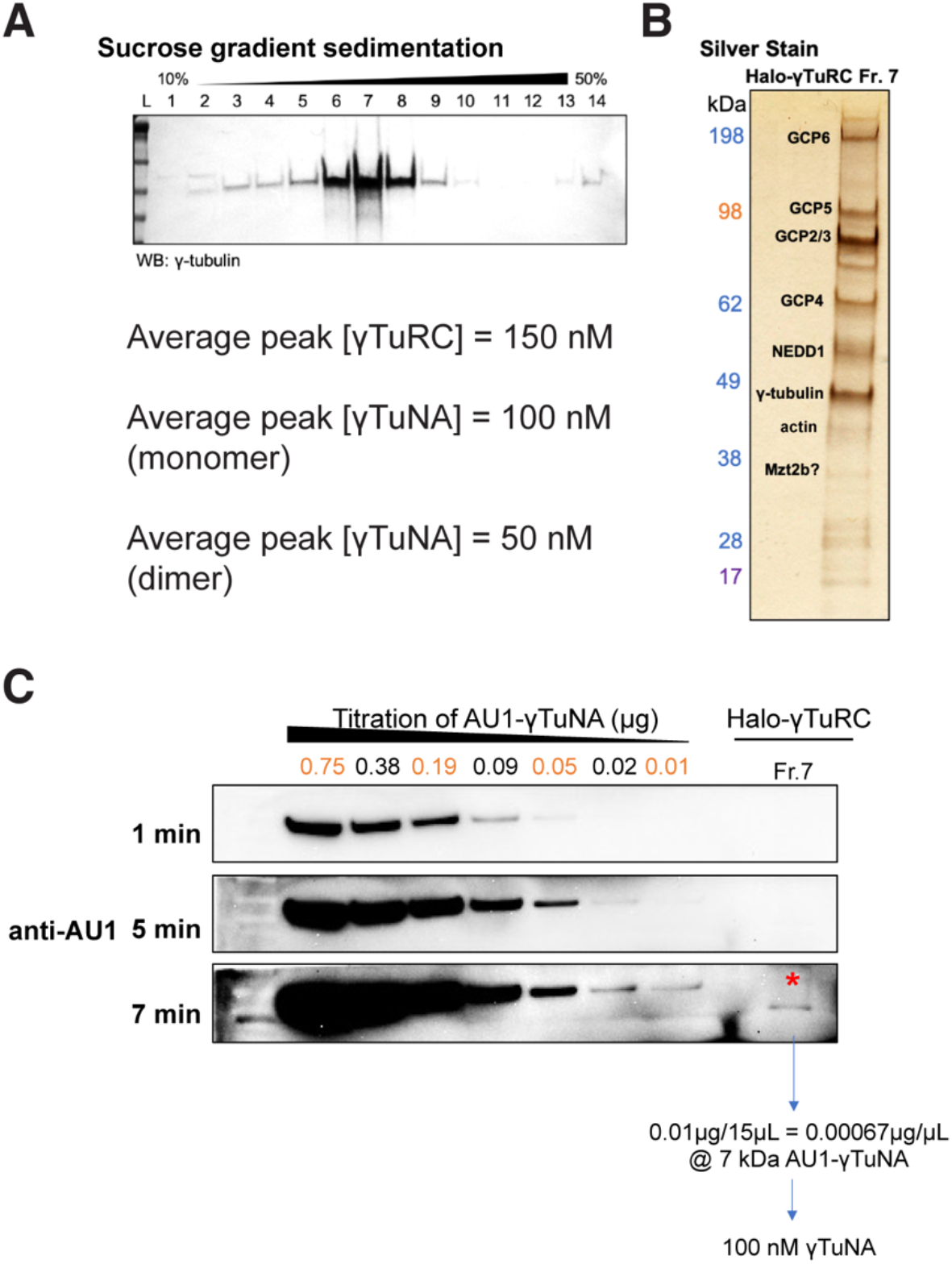
Purity and concentration assessment of γTuRCs purified via Halo-γTuNA pulldown. **A**) Western blot of γ-tubulin in fractions from a 10-50% w/w sucrose gradient. The γTuRC peak was routinely located in fraction 7, as shown in A. **B)** SDS-PAGE and silver stain of the peak γTuRC fraction. Dominant bands are known γTuRC components, as verified by mass spectrometry. **C**) Western blot-based determination of γTuNA concentration remaining in γTuRC peak fraction. For this, Halo-γTuRC preps were done with Halo-3C-AU1 reporter tagged versions of human γTuNA. After elution of target protein via PreScission (3C) cleavage, any remaining γTuNA will still be fused to the AU1 epitope. Probing of the γTuRC peak fraction with a sensitive anti-AU1 antibody (against a titration of known amounts of AU1) allowed us to determine that 100 nM of AU1 remained. This corresponded to 50 nM γTuNA dimer.

**Figure S4.**
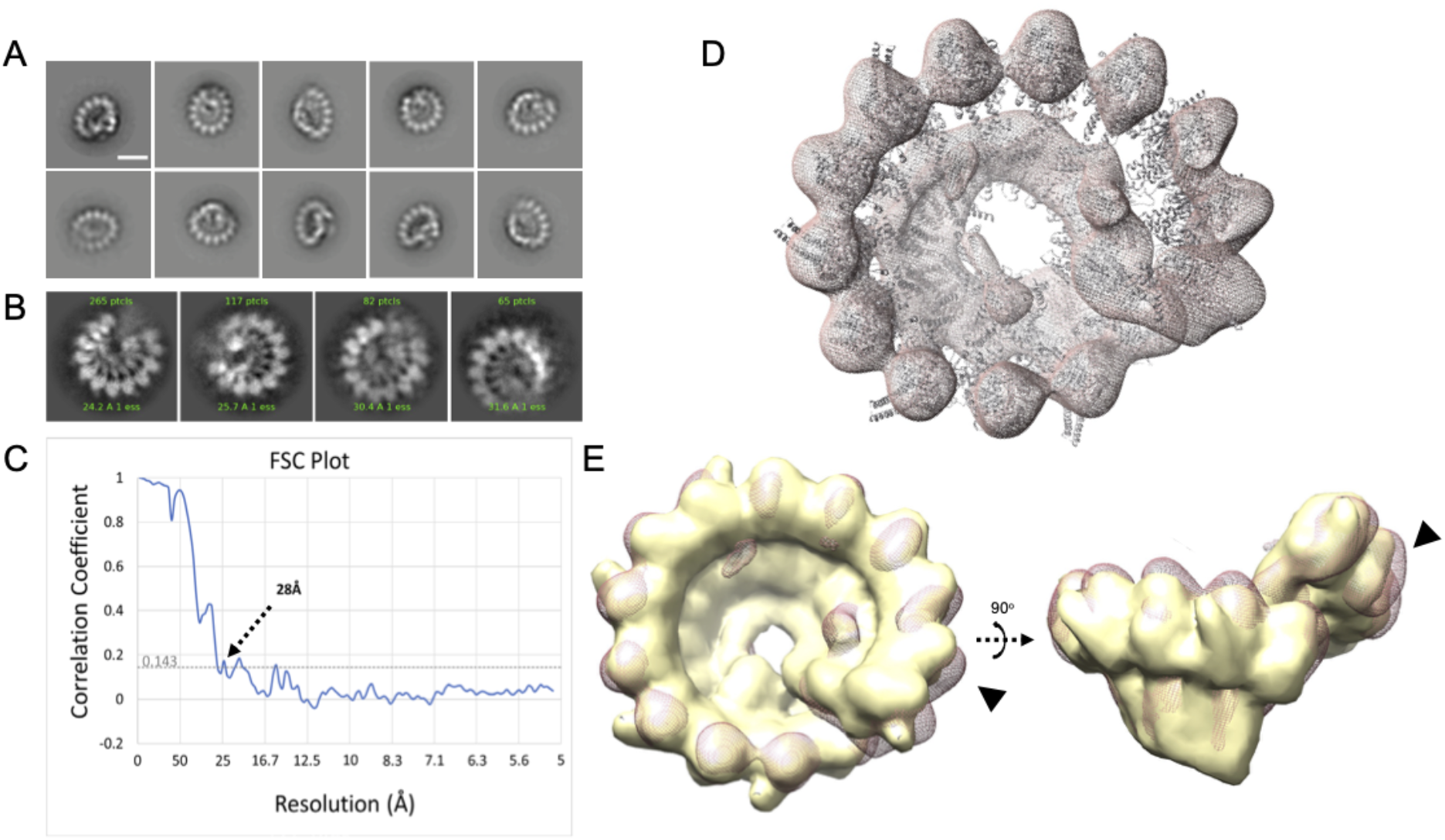
γTuRCs purified via Halo-γTuNA pulldown are fully assembled rings. Electron-microscopy data of γTuNA-prepped γTuRC. **A)** 2D class averages from 2,692 particles isolated from negative-stain data. Scale bar = 20 nm. **B)** Selected 2D class averages from 529 particles selected from cryoEM data. **C**) Fourier shell correlation (FSC) plot from 3D reconstructions and final refinements of particles of γTuRC from negative-stain EM data, showing a final resolution of 28 Å. **D**) Map of γTuNA-prepped γTuRC (red; mesh) with previously published holo-γTuRC structure from *Xenopus laevis* docked. **E**) Overlay of γTuNA-prepped γTuRC with a 28Å simulated map from the published *Xenopus laevis* structure of γTuRC (yellow), showing a high correlation between the two (0.89). Highlighted is the area where γTuNA is predicted to bind γTuRC (arrows). Figure made using Chimera. **D and E**) Both parts use structure PDB 6TF9 for comparison.

**Figure S5.**
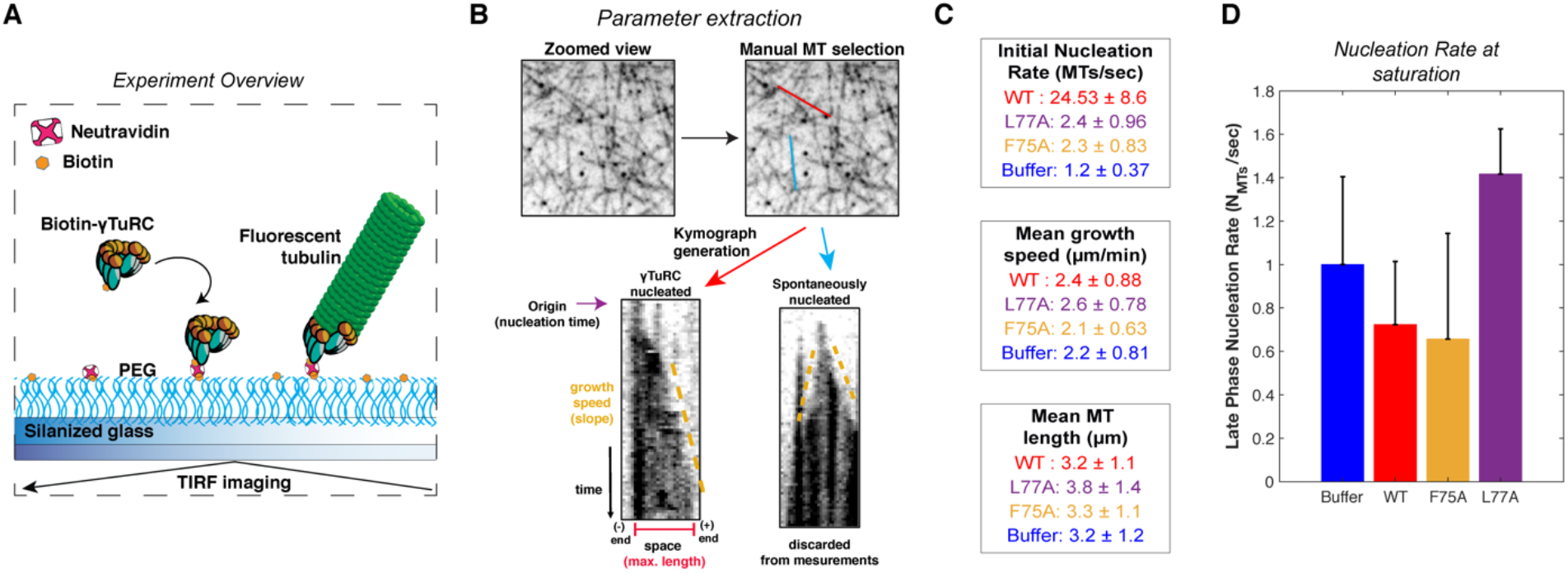
Overview and additional single molecule TIRF data. **A**) Diagram of single molecule TIRF assay for γTuRC MT nucleation. **B**) Process diagram of parameter extraction from single molecule TIRF data: individual MTs were manually selected in FIJI (ImageJ, NIH). Next, time-lapse data (stack) was resliced for each MT to generate space vs time plots (better known as kymographs). From these kymographs, we determined whether a MT nucleated from a γTuRC (no growth at one end) or spontaneously (growth from both ends). If a MT was nucleated by a γTuRC, then we manually extracted the time that MT nucleated (origin), the growth speed, and the max length of that MT in the kymograph. If spontaneously nucleated, we did not extract these parameters. **C**) Numerical representations of data presented in Fig. 6C-F. Mean ± standard deviation. **D**) Nucleation rate measured at saturation (30 to 150 s), normalized to the buffer mean. Error bars are standard deviation.

**Figure S6.**
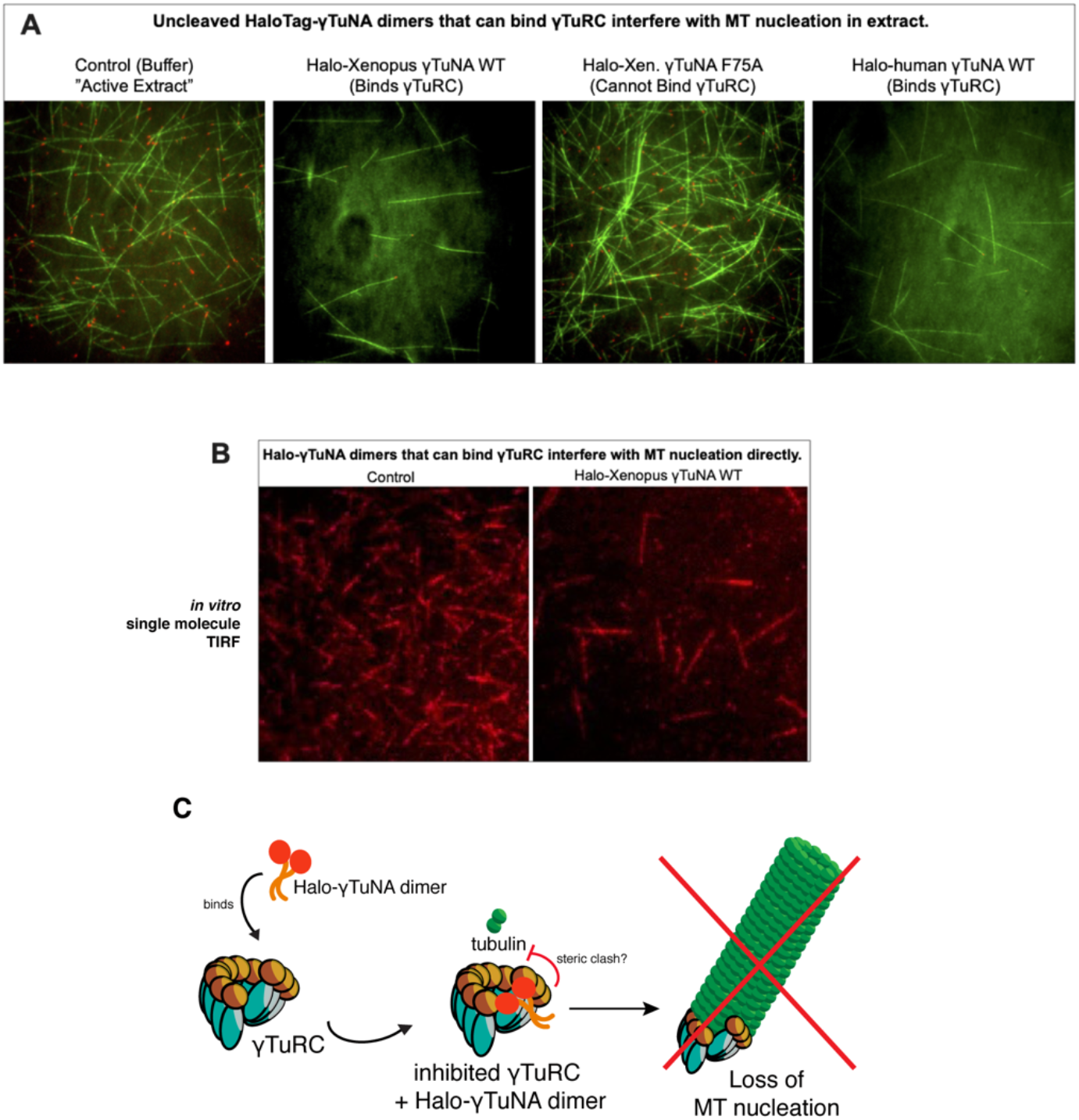
The presence of large, bulky N-terminal tags on γTuNA directly inhibits γTuRC activity in extract and in vitro. **A**) TIRF assay of *Xenopus* egg extract in the presence of N-terminally tagged, Halo-γTuNA constructs (wildtype and F75A) after 15 min. Alexa488-tubulin and EB1-mCherry were used to visualize MTs and MT plus-ends. Halo-γTuNA constructs that can bind γTuRC (as determined in Fig. 3) drastically decreased the amount of MTs nucleated (human and *Xenopus* wildtype). Halo-γTuNA F75A mutant, which cannot bind γTuRC, had no effect on MT levels. Images were taken at 18-20°C. Halo-γTuNA was added at 2 μM final. **B**) Single molecule TIRF assay of γTuRC-mediated MT nucleation *in vitro*. After 5 min, γTuRCs alone efficiently nucleate MTs. In the presence of 3.3 μM wildtype *Xenopus* Halo-γTuNA, few γTuRCs nucleate MTs, indicating a direct inhibition of γTuRC activity. **C**) Model for inhibition of γTuRC activity by N-terminal Halo-tagged γTuNA dimers.

**Figure S7.**
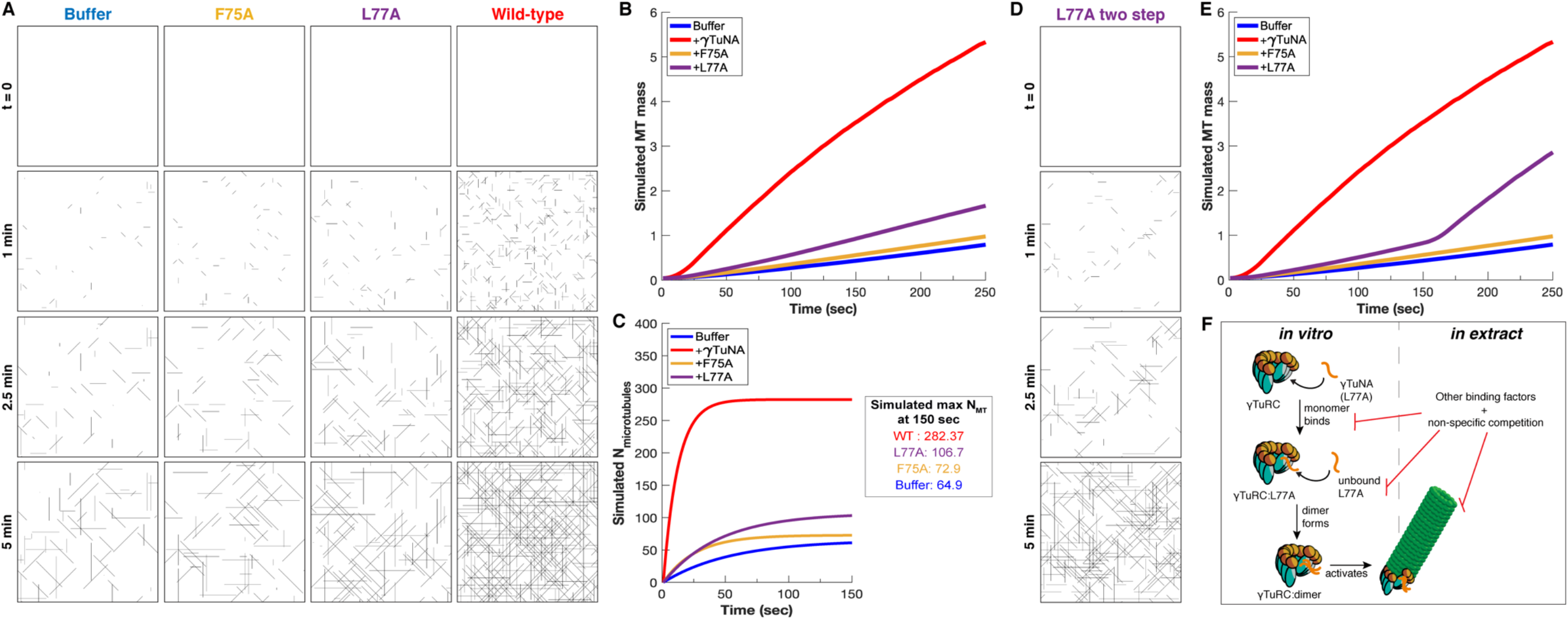
Simulation of γTuNA’s effect on γTuRC MT nucleation activity. **A)** Simulated frames of MT nucleation and growth (simulated 40 x 40 μm^2^ plane) over 5 minutes. Based on just the initial nucleation rate, the simulated data correlates well with the observed data in Fig. 6 (see Video-3 for side-by-side comparison). **B)** Simulated MT mass over time, measured from simulated movies using the same FIJI (ImageJ) pipeline as used with real data in Fig. 6. **C)** Simulated number of nucleated MTs over time, with simulated maximum number of MTs at 150s. **D)** Simulated frames of MT nucleation and growth of a two-step model for the L77A mutant. **E)** Simulated MT mass over time for the two-step L77A model, now reflecting a late stage increase in MT mass, similar to what was observed in Fig. 6B. **F)** Model for L77A’s two stage behavior: involving sequential binding of two separate L77A monomers that form a dimer on γTuRC before triggering its activation. In extract, this behavior is not observed, suggesting the presence of other extract factors compete against this interaction.

## Notes

### Competing Interest Statement

The authors have declared no competing interest.

