## Supplementary material for "The conserved centrosomal motif, γTuNA, forms a dimer that directly activates microtubule nucleation by the γ-tubulin ring complex (γTuRC)": MATLAB-Numerical Simulation of gTuNA-gTuRC MT nucleation

%% Numerical simulation of MT mass over time using initial nucleation rates and mean growth speed.
% by Michael Rale, Petry Lab, Princeton University
%

%Requires Matlab R2019a or later. Can also use GNU-Octave.
%From single molecule TIRF assays, the nucleation rate (# of MTs nucleated per sec) and growth rate (microns/min)
%are used to simulate gTuRC mediated MT nucleation in the presence or
%absence of different gTuNA constructs (or buffer).

%Assuming the mass contribution of just nucleated MTs is negligible in their starting time step:
%Assuming no catastrophes:

clear
clc
close all

%Experimentally determined values.
MeanSpeed = 2.3593;             %experimentally measured, averaged, in micron per minute.
Buffer = [1.2401; 66.2193];     %Mean initial nucleation rate in MTs/sec; average max number of MTs observed in each rxn.
Wildtype = [24.5338; 293.0516]; %Mean initial nucleation rate in MTs/sec; average max number of MTs observed in each rxn.
F75A = [2.2621; 72.8764];       %Mean initial nucleation rate in MTs/sec; average max number of MTs observed in each rxn.
L77A = [2.4351; 107.9898];      %Mean initial nucleation rate in MTs/sec; average max number of MTs observed in each rxn.

conditions = [Buffer Wildtype F75A L77A L77A]; 

%Initialize simultation with constants.
growthSpeed = MeanSpeed/60;     %per second
totalTime = 300;                %in seconds
Sim = [];
[~, experiments] = size(conditions);

%Run numerical simulation.
for j = 1:experiments
    %Variables that need to be reset each time.
    MTmass = 0;
    MTnumber = 0;
    NewMT = 0;
  
    %Simulated MT nucleation, growth
    for i = 0:totalTime+1
        
        %Determine new MT mass added each step.
        MTmass = MTmass + (MTnumber * growthSpeed);
        
        %Nucleate new MTs according to nucleation rate (with special case
        %for L77A late-stage behavior)
        if j == 5 && i > 150
              NewMT = conditions(1,2) * (1 - (MTnumber)/( conditions(2,2) - tempSim(150,2) ) ); %constants from WT parameters for L77A late-stage
        elseif i > 0
              NewMT = conditions(1,j) * (1 - (MTnumber/conditions(2,j))); % see Equation #2 in Methods.     
        end
        
        %Determine total MT numbers nucleated up to this time step.
        MTnumber = MTnumber + NewMT;

        %Temporary storage of simulated data.
        tempSim(i+1, 1) = i;
        tempSim(i+1, 2) = MTnumber;
        tempSim(i+1, 3) = MTmass;
    end
    
    %Aggregate simulated data from all experiments.
    Sim = [Sim tempSim];
end

%Remove 0th timepoint.
Sim = Sim(2:end, :);

%Split data according to condition to pass on to visual simulation script.
BufferSim(:,1:3) = Sim(:, 1:3);
WTSim(:,1:3) = Sim(:, 4:6);
F75ASim(:,1:3) = Sim(:, 7:9);
L77ASim(:,1:3) = Sim(:, 10:12);
L77ASim2(:,1:3) = Sim(:, 13:15);

%Save this output as a .mat file, along with growthSpeed variable for use
%in the visual simulation.

 save('Rale_NumericalMTsimulation.mat', 'BufferSim', 'WTSim', 'F75ASim',...
     'L77ASim', 'L77ASim2', 'growthSpeed');

 
% % % END % % %
