## Supplementary material for "The conserved centrosomal motif, γTuNA, forms a dimer that directly activates microtubule nucleation by the γ-tubulin ring complex (γTuRC)": MATLAB- Graphical Simulation of gTuNA-gTuRC MT nucleation

%% Visual simulation of microtubule nucleation by gTuRC (in vitro, single molecule TIRF assay)
% by Michael Rale, Petry Lab, Princeton University
%

%Requires Matlab R2019a or later.
%From single molecule TIRF assays, the nucleation rate (# of MTs nucleated per sec) and growth rate (microns/min)
%are used to simulate a field of MTs nucleating and growing under different
%conditions.

% Needs the NumericalMTsimulation.mat file that 'Rale_NumericalMTSimulation.m' file generates.
% Make sure it is in the same Current Folder as this script.

%This simulation will display the frames as they are generated.  You can
%just view them without saving, or save them, by selecting the option when
%it shows up in the command window.

%% First, generate single MTs growing over time and store them.
...Then the stored MTs will be displayed together later.
    
clear
clc
close all

%Load in the numerical simulation data based on initial nucleation rate.
load('Rale_NumericalMTsimulation.mat')
condition = [BufferSim(:,2) WTSim(:,2) F75ASim(:,2) L77ASim(:,2) L77ASim2(:,2)];  %refers to the column vector with # of MTs at each time.
prompt1 = 'Pick a condition: \n\n Buffer = 1, WT = 2, F75A = 3, L77A = 4, L77A with late-stage WT binding = 5 \n\n';
option = input(prompt1);  
prompt2 = 'Do you want me to save video frames of the simulation? \n\n Yes = 1, No = 0 \n\n';
option2 = input(prompt2);  

%Constants for simulation
growthspeed = growthSpeed;                  % micron per sec. Imported from the numerical simulation.
steps = 300;                                % iterations of the same MT (AKA the time); one step in this simulation will equal 1 second of real.
boxLorH = 40;                               % set length and width in microns (looking for square view, so both must be the same).


%Convert sim data into explicit number of new MTs at each step.
[row, ~] = size(condition(:,option));
newMTsPerStep = zeros(row,1);
for k = 1:row
    if k == 1
        newMTsPerStep(k, 1) = round(condition(1,option));
    else
        newMTsPerStep(k, 1) = round(condition(k, option)) - round(condition(k-1, option));
    end
end


%Total Number of simulated MTs (for sanity check later).
numMTs = sum(newMTsPerStep(1:steps,1)); 

%Set up some stored random values for determining initial position and
%orientiation of MTs. Three levels of randomization.
rng('shuffle');                         %initialize the random number generator. Needs to be 'shuffle' to make sure that each time this is run the placement of MTs is different.   
rand_x = randi([0 boxLorH], 1, 1000);
rand_y = randi([0 boxLorH], 1, 1000);
dir1 = randi([1 8], 1, 1000);           %to generate a large list of randomly determined initial directions of each MT.

% Initialize a place to store all MTs that are about to be generated.
AllMTs_x = zeros(numMTs, steps+2);
AllMTs_y = zeros(numMTs, steps+2);

% Generate a new MT and assign it with its direction, calculate the new plus-end.  
for j = 1:numMTs                        % Each new j is a new microtubule.

%Randomly pick an origin for the MT.
origin = [rand_x(randi(1000)) rand_y(randi(1000))];   %stored as [x y] notation

%Pick a direction to grow in.
direction = (dir1(randi(1000)));  %should result in a single integer between 1 and 8.

%Initialize the plus-end and variables.
plusend = origin;                 %x, y location of plus end per row, at the start of each MT's life, is the origin
delayID = [];
windows =zeros(steps, 2);

%Based on that direction, calculate the new plus-end after growth.
for i = 1:steps
    if direction == 1
        newplusend = [plusend(i,1)+growthspeed plusend(i,2)];  %case 1, or 0deg or 360deg (x gets all the growth, y gets nothing)
    elseif direction == 2
        newplusend = [plusend(i,1)+(growthspeed/sqrt(2)) plusend(i,2)+(growthspeed/sqrt(2))]; %for case 2, or 45 degrees.
    elseif direction == 3
        newplusend = [plusend(i,1) plusend(i,2)+growthspeed];  %case 3, or 90deg (x gets nothing, y gets growth)
    elseif direction == 4
        newplusend = [plusend(i,1)-(growthspeed/sqrt(2)) plusend(i,2)+(growthspeed/sqrt(2))]; %for case 4, or 135deg.
    elseif direction == 5
        newplusend = [plusend(i,1)-growthspeed plusend(i,2)];  %case 5, 180deg (x gets all the growth, y gets nothing)
    elseif direction == 6
        newplusend = [plusend(i,1)-(growthspeed/sqrt(2)) plusend(i,2)-(growthspeed/sqrt(2))]; %for case 6, or 225deg.
    elseif direction == 7
        newplusend = [plusend(i,1) plusend(i,2)-growthspeed];  %case 7, or 270deg (x gets nothing, y gets growth)
    elseif direction == 8
        newplusend = [plusend(i,1)+(growthspeed/sqrt(2)) plusend(i,2)-(growthspeed/sqrt(2))]; %for case 8, or 315deg.
    end
%     MTlength(end+1,1) = MTlength(end,1) + growthspeed; %MT length not
%     being used right now
    plusend = cat(1,plusend,newplusend);
    
    leftwindow = (sum(newMTsPerStep(1:i)));
    rightwindow =(sum(newMTsPerStep(1:(i+1))));
    windows(i,:) = [leftwindow rightwindow];
    
    % Record the step in which each MT will first appear.
    if i == 1
       delayID(1:newMTsPerStep(1,1),1) = i;
    elseif (i > 1) && (i < steps - 1)
       delayID(windows(i-1,1):windows(i,1), 1) = i;
    end
end

[row, ~] = size(plusend);


MT_x = (origin(1));
MT_y = (origin(2));

%Generate a single MT growing over time.
for i = 1:row   
   %MT coordinates for this frame and store.
    MT_x = [MT_x, plusend(i,1)];
    MT_y = [MT_y, plusend(i,2)];
%     MT_x(1, i+1) = plusend(i,1);  %For pre-allocation, but I didn't notice a speed increase.
%     MT_y(1, i+1) = plusend(i,2);  %For pre-allocation, but I didn't notice a speed increase.
end

%Store this MT
    AllMTs_x(j,:) = MT_x;
    AllMTs_y(j,:) = MT_y;
    
%clean up windows
close all
end

%Add each new MT according to the nucleation rate for each condition.

delayedMTs_x = zeros(size(AllMTs_x));   %preallocation
delayedMTs_y = zeros(size(AllMTs_x));   %preallocation
frontRow = [];
backRow = [];

for i = 1:numMTs
    frontRow(1,:) =  repelem( AllMTs_x(i,delayID(i,1)) , delayID(i,1) );
    backRow(1,:) = AllMTs_x(i,((delayID(i,1)+1):end));
    x = horzcat(frontRow(1,:),backRow(1,:));
    delayedMTs_x(i, 1:length(x)) = x(1, 1:length(x));
    frontRow = [];
    backRow = [];
end

for i = 1:numMTs
    frontRow(1,:) =  repelem( AllMTs_y(i,delayID(i,1)) , delayID(i,1) );
    backRow(1,:) = AllMTs_y(i,((delayID(i,1)+1):end));
    x = horzcat(frontRow(1,:),backRow(1,:));
    delayedMTs_y(i, 1:length(x)) = x(1, 1:length(x));
    frontRow = [];
    backRow = [];
end


%% Now generate the simulated video frames.
close all

%Sets up folders to save plot frames. These output frames can be turned into a single movie using
%Quicktime Player-->"Open image sequence"--->save converted mov.
%Or use ImageJ's import-->image sequence option --> select these frames-->save as TIFF or AVI.

labelcondition = {'BufferVid', 'WTVid', 'F75AVid', 'L77Asim1Vid', 'L77Asim2Vid'};

%Check if a folder already exists for this condition.
%If so, remove that folder and replace with this new one.
%If not, make a folder to store the frames.

if isfolder(string(labelcondition(option)))
    rmdir(string(labelcondition(option)))
    mkdir(string(labelcondition(option)))
else
    mkdir(string(labelcondition(option)))
end

%Move to the newly created folder and save frames as .PNG format.
cd(string(labelcondition(option)))
format = 'png';

[row, col] = size(delayedMTs_x);

%Start the figure and plotting for the simulation.
figure
    %Appearance settings.
    box off
    xlim([0 boxLorH])
    ylim([0 boxLorH])
    set(gcf,'position',[300,300,20*boxLorH,20*boxLorH])
    set(gca,'xtick',[])
    set(gca,'ytick',[])
    hold on
    
    %Plot all MTs.
for i = 1:col %each frame
    for j = 1:row %update new plus-end location for every microtubule belonging to this step.
        if ~((delayedMTs_x(j,1) == delayedMTs_x(j,i)) && (delayedMTs_y(j,1) == delayedMTs_y(j,i)))  % having this here lowers the run time by 20% on average.
            plot([delayedMTs_x(j,1) delayedMTs_x(j,i)], [delayedMTs_y(j,1) delayedMTs_y(j,i)], 'LineWidth',1, 'color', 'k' )
        end
    end
    
    pause(0.05)  %this pause is here to allow the user to see the plot and allow time for the computer to properly save the frame.
    
    %If user wants to save the frames from the simulation, it will do so.
    if option2 == 1
        %grab only the figure with no border or white space around it.
        F = getframe(gca);  
        %save the frame to the folder.
        imwrite(F.cdata, string(i)+"_frame_"+string(labelcondition(option))+"."+string(format));   
    end
end
    hold off
close all


%Return the directory to the original location.
cd ../

%Tell user that sim has finished.
disp('Simulation for the chosen condition has finished.');

% % % END % % %
